# cWnt signaling regulates the early proliferative response and the differentiation of the neural, muscle and parenchymal tissues during planarian whole-body regeneration

**DOI:** 10.64898/2026.11.12.766096

**Authors:** P. Coronel-Córdoba, J. Gaya-Ocón, H. García-Castro, E. Emili, N. J. Kenny, S. Chitgaran, G. Tapia-Saló, J.F. Abril, E. Saló, J. Solana, T. Adell

## Abstract

Organizers, or signaling centers, play a central role in coordinating tissue growth, patterning, and differentiation during development and regeneration. In particular, organizers that arise during the regeneration of specific animal structures typically involve components of the canonical Wnt (cWnt) pathway and contribute to tissue growth and differentiation. However, how organizers and the cWnt pathway promote these processes in animals capable of whole-body regeneration remains poorly explored. Here, we analyze organizer functions in planarians, which can regenerate any missing body part from a population of adult pluripotent stem cells. The tips of the planarian body have been proposed to act as organizers. Here we demonstrate the organizing capacity of these tips by showing that anterior or posterior tips grafted into a host animal induce the formation of a patterned head or tail, respectively. Given that the cWnt inhibitor Notum and the activator Wnt1 define these tip regions, we next investigate the role of cWnt signaling in promoting growth and cell differentiation. Our results reveal that cWnt inhibition anteriorly and activation posteriorly are required to promote stem cell proliferation during regeneration. Furthermore, we show that repression and activation of cWnt signaling through *βcatenin-1* and *apc* silencing, respectively, can rescue regeneration in sublethally irradiated planarians. Moreover, we show how cWnt signaling at posterior wounds is necessary for the differentiation of specific cell types, including neurons, muscle, and parenchymal cells. Together, our findings identify cWnt signaling as a key regulator of growth and cell differentiation within planarian organizing regions and provide new insight into how organizers control whole-body regeneration.

## Introduction

How growth, patterning, and differentiation are orchestrated to rebuild lost structures is a central question in regenerative biology. Key to this coordination are organizers, also called organizing or signaling centers: groups of cells that, through the secretion of morphogens, integrate the growth, patterning, and differentiation of the surrounding tissues during both embryogenesis and regeneration^1–5^. The first organizers to be identified were the head organizer of *Hydra*, which patterns the oral-aboral axis^6^, and the Spemann-Mangold organizer of the salamander gastrula^7^, which patterns the dorsoventral and antero-posterior axes while promoting the proliferation and differentiation of the neural tube, somites, and notochord^8^. Other organizers act more locally, such as the isthmic organizer at the midbrain-hindbrain boundary of vertebrate embryos, where Wnts and FGFs promote the proliferation and differentiation of neural progenitors^9^. The vertebrate limb is a particularly informative example of tissue growth under the control of an organizer, as it allows embryonic development and regeneration to be compared directly. During embryogenesis, the apical ectodermal ridge (AER) of the limb bud secretes FGFs that induce the proliferation of mesenchymal cells^10^, whereas during salamander limb regeneration the apical epithelial cap (AEC), which also secretes FGFs and Wnts, maintains the proliferation of blastema cells while controlling their differentiation into muscle, bone, neurons, and epidermal cells^11^. Beyond the limb, organizers are known to play an important role in promoting regeneration in animals that regenerate specific structures. During both Xenopus tail and zebrafish fin regeneration, for instance, an organizing center expressing Wnts and other morphogens is essential to drive the growth of the regenerating structure^12,13^. Likewise, in mammalian digit tip regeneration, the nail epithelium displays an organizer-like function, as Wnt activation in this tissue is required to attract the nerves that promote the growth of the forming blastema^14^.

Organizing centers also play a central role in animals capable of whole-body regeneration, such as *Hydra* and planarians. In *Hydra*, the head organizer expresses *wnt3* and *brachyury* and is known to promote oral identity during regeneration^15–17^. In planarians, the head and tail tips have been proposed to function as organizers^18,19^, potentially acting in a manner similar to the organizer of adult *Hydra*. The signals that define planarian organizers are morphogens of the canonical Wnt (cWnt) pathway. At the anterior and posterior body tips, at the intersection between the dorsoventral boundary (DVb) and the midline, a group of muscle cells expresses the Wnt inhibitor *notum*^20^, in the anterior tip, and *wnt1*^21^ in the posterior. RNAi-mediated inhibition of *notum* and *wnt1* blocks head and tail regeneration, respectively, producing headless and tailless animals^20,22^. The most extreme phenotype after *notum* or *wnt1* inhibition is a polarity shift that generates two-tailed or two-headed animals, respectively, demonstrating the essential role of these morphogens in axial patterning, as previously described in *Hydra*^23^.

Therefore, in animals capable of whole-body regeneration, the function of organizers has been linked mainly to axial patterning, whereas in animals that regenerate specific structures their influence extends to tissue growth and differentiation. Planarians offer an ideal system to ask whether organizers also regulate tissue growth and differentiation in the context of whole-body regeneration^24^. Planarian regenerative capacity relies on an adult population of pluripotent stem cells, the neoblasts, which proliferate and differentiate upon amputation to regenerate any lost structure^25^. Thus, planarians are a powerful model to study cell proliferation, tissue differentiation, and morphogenesis in general, since their complex tissues must be correctly regenerated and integrated with the pre-existing ones^26^. These tissues include the body-wall muscle, composed of circular, diagonal, and longitudinal fibers that are responsible both for motor function and for the expression of morphogens and other position control genes (PCGs)^27,28^. Planarians also present a centralized bilateral nervous system connected to a pair of eyes^29^, a monolayered epidermis^30^, and a branched gut composed of basal cells, goblet cells, and phagocytes^31^, which connects to a pharynx that serves as both mouth and anus^32,33^. Finally, planarians have a heterogeneous population of cells referred to as parenchymal, which includes glia and pigment cells, that forms a supporting tissue in which other cell types are embedded^34–36^, such as secretory cells, neoblasts, and protonephridia^37^.

As in the pioneering grafting experiments performed by Ethel Browne in *Hydra*^6^, transplantation has also been used in planarians to induce the growth of new axes and to test the organizing activity of different regions. As exposed, the anterior and posterior organizers, defined by the expression of *notum* and *wnt1*, respectively, are located at the tips, where the DVb and the midline intersect. In turn, the DVb and the midline are defined by members of the TGF-β pathway; *bmp*, expressed in the dorsal midline^38^, and *admp*, expressed in the ventral midline and at the DVb^39^, which acts together with *bmp* in a dorso-ventral regulatory circuit. Grafting dorso-ventrally inverted fragments into a host induces the formation of a complete Siamese planarian, demonstrating that the confrontation of dorsal and ventral signals is sufficient to trigger growth^40^. This observation is consistent with the events that occur during normal regeneration, in which dorsal and ventral tissues are brought into contact upon wound closure through an immediate muscle contraction^27,41^. The requirement for dorso-ventral confrontation to promote regeneration is further supported by the phenotypes observed after inhibition of BMP/ADMP pathway components, which impairs the formation of a proper DVb and blocks regeneration^38,39,42^. More recently, it has been shown that grafting of the DVb itself is sufficient to induce an outgrowth, and that inhibition of βcatenin-1, the cWnt pathway effector, only confers anterior identity in the DVb region, turning it into a head-like structure^43^. This leaves open the question of whether cWnt signaling is required for processes beyond axial patterning. However, owing to their technical difficulty involved, few studies have succeeded in grafting the tips themselves, which express *wnt1* and *notum* and are therefore expected to hold organizer activity^1^. The only published result shows that grafting the anterior tip can induce the formation of an anteriorly polarized outgrowth, which nevertheless lacks an organized midline and dorso-ventral polarity^44^. Thus, it remains unknown whether grafting the anterior tip can induce a patterned head, or whether grafting the posterior tip can induce any kind of growth. Moreover, it is not known whether the cWnt signals present at the tips are likewise restricted to patterning the outgrowth or whether they also promote the growth and differentiation of the forming tissue. This question is of major importance because, although cWnt signaling has been linked to tissue growth and differentiation in several embryonic and regenerative contexts, its role in these processes during whole-body regeneration remains far less explored.

To address these questions, we first demonstrate that grafting the anterior and posterior planarian tips induces the formation of a patterned head and tail, respectively. To determine whether the planarian tips are required not only for identity specification but also for promoting tissue growth and differentiation, two hallmark features of organizing regions, we then examined whether modulation of cWnt signaling promotes proliferation and affects cell differentiation during planarian regeneration. Our results show that inhibition of cWnt signaling anteriorly and its activation posteriorly promotes neoblast proliferation and can rescue regeneration in sublethally irradiated planarians. Moreover, we show that cWnt signaling is necessary during tail regeneration for the differentiation of specific planarian tissues, including ChAT neurons, parenchymal cells, and the body-wall muscle.

## Results

### Grafting of the anterior and posterior tips can induce the formation of a fully patterned head and tail

The anterior and posterior tips of adult planarians, which express the Wnt inhibitor *notum* (**Fig 1A**) and *wnt1* (**Fig 1B**), respectively, have been proposed to act as signaling centers or organizers^1^. A previous study showed that grafting the anterior tip induces either an outgrowth without apparent A-P organization (“Tubular”, **Fig 1C**) or an outgrowth with anterior identity but lacking an organized midline and dorso-ventral polarity (“Polarized”, **Fig 1D**)^44^. However, we reasoned that, if the planarian tips have organizing capabilities, their grafting should be able to fully pattern the growing structure (“Bilateral”, **Fig 1E**). To test this, we grafted the planarian tips following a published protocol^45^ (**Fig 1F and S1A Fig**). When the anterior tip of a donor animal was grafted into the post-pharyngeal region of a host (**Fig 1G**), 8 of the 15 successfully integrated grafts developed the previously reported “Polarized” outgrowth^44^ (**Fig 1F and S1B-C Fig**). These outgrowths differentiated one eye on each side (**S1C Fig**), and accumulated neural cells distally without a bilateral brain organization (**S1D Fig**). In contrast, 6/15 grafts produced a patterned head, “Bilateral” outgrowth, with two bilaterally arranged eyes as in a normal planarian head (**Fig 1H-H’**), a well-defined DVb (**Fig 1I-I’**), and a bilobed brain connected by a commissure, similar to that of normal heads (**Fig 1J-J’**). Grafting the posterior tip into the pre-pharyngeal region (**Fig 1K**) produced, in 9/15 cases, outgrowths resembling those described for anterior tip grafts^44^ (**Fig 1F and S1E-F Fig**), but lacking eyes and containing only scattered neural cells (*pc2*^+^ cells) (“Polarized-like”, **S1F-G Fig**). Importantly, 3/15 grafts induced a patterned tail (“Bilateral-like”, **Fig 1L-L’**), with a defined DVb (**Fig 1M-M’**) and two structures visible with nuclear DAPI staining and surrounded by *pc2*^+^ neural cells, resembling the posterior ventral nerve cords (VNCs) of normal tails (**Fig 1N-N’**). The remaining 3/15 posterior grafts produced tubular outgrowths (**Fig 1F**). Finally, to test whether the signals of the DVb or the midline alone could induce a patterned outgrowth, we grafted the DVb region from the head and the midline from the pre-pharyngeal region into the post-pharyngeal region of a host. DVb grafts promoted only “Tubular” outgrowths (**Fig 1F and 1O**), as previously described^40^, while midline grafts never promoted an outgrowth (**Fig 1F and 1P**).

**Figure 1.**
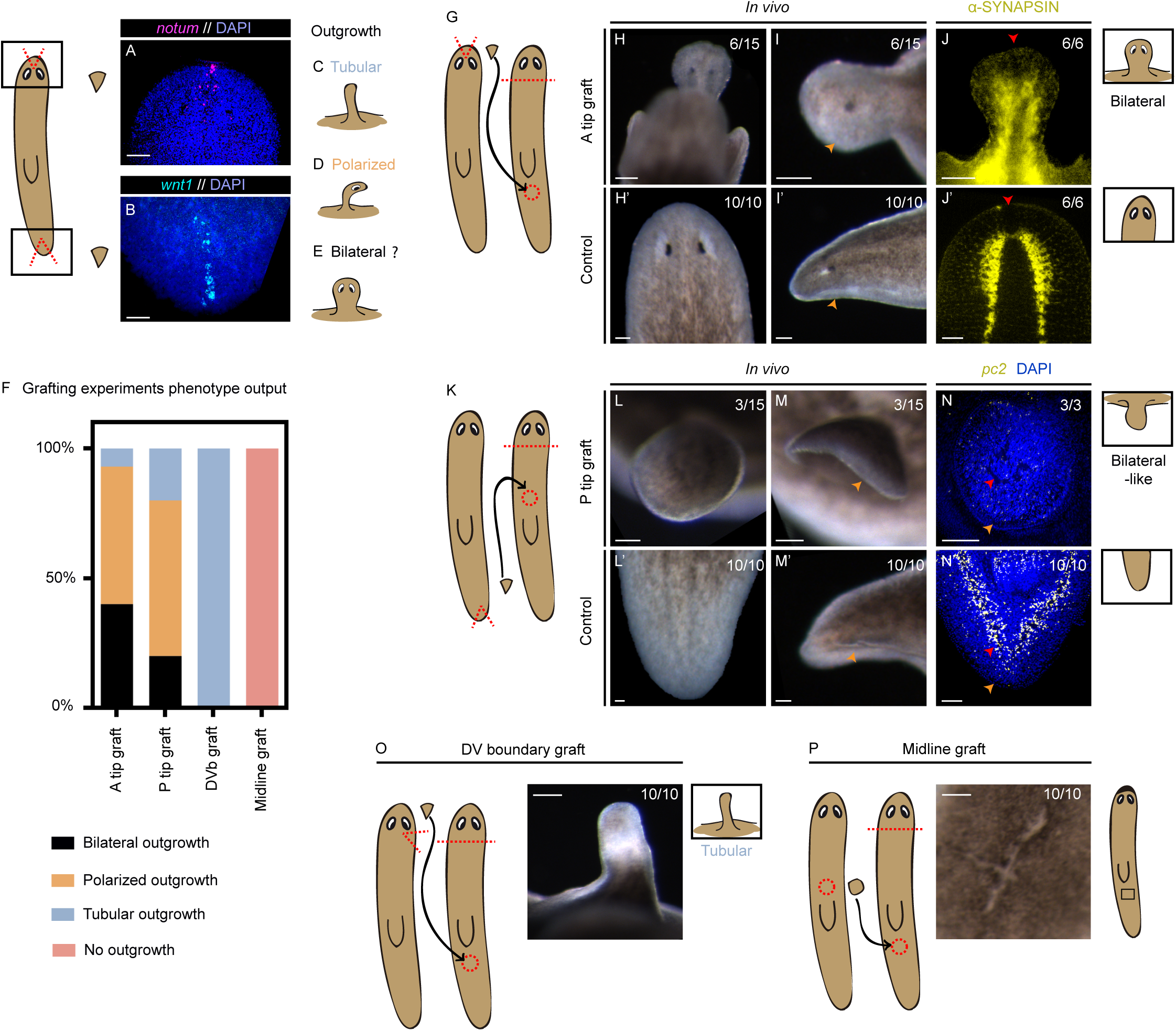
Grafting of the anterior and posterior tips can induce the formation of a patterned head and tail. A-B. FISH showing *notum* (magenta) in an intact planarian head (A) and *wnt1* (cyan) in an intact planarian tail (B). Nuclei are labeled with DAPI (blue). **C-E.** Schematic representation of the possible outcomes after grafting: Tubular outgrowth without patterning (C), Polarized outgrowth with A-P polarity (D) and Bilateral outgrowth with A-P and D-V polarity (E). **F.** Stacked bar plot showing the proportion of each outcome after grafting the anterior tip, the posterior tip, the DVb beneath the eyes or a piece of the pre-pharyngeal midline. **G.** Schematic representation of the anterior tip graft into the post-pharyngeal region of the host. **H-I’.** *In vivo* dorsal (H, H’) and lateral (I, I’) views of an anterior outgrowth 5 weeks after grafting (H, I) and of a control head 5 weeks after amputation (H’, I’). Orange arrows indicate the DVb. **J-J’.** α-synapsin immunostaining (yellow) of an anterior outgrowth 5 weeks after grafting (J) and of a control head 5 weeks after amputation (J’). Red arrows indicate the brain commissure at the midline. **K.** Schematic representation of the posterior tip graft into the pre-pharyngeal region of the host. **L-M’.** *In vivo* dorsal (L, L’) and lateral (M, M’) views of a posterior outgrowth 5 weeks after grafting (L, M) and of a control tail 5 weeks after amputation (L’, M’). Orange arrows indicate the DVb. **N-N’.** FISH showing *pc2* (yellow) in a posterior outgrowth 5 weeks after grafting (N) and in a control tail 5 weeks after amputation (N’). Red arrows indicate the VNC-like structures and orange arrows indicate the DVb, both visible with DAPI nuclear staining (blue). **O.** DVb graft from the region beneath the eyes of the donor into the post-pharyngeal region of the host, resulting in a tubular, unpatterned outgrowth. **P.** Midline graft from the pre-pharyngeal midline (between the VNCs) of the donor into the post-pharyngeal region of the host, showing wound closure without outgrowth formation. DVb: dorsal-ventral boundary. FISH: whole-mount fluorescent *in situ* hybridization. VNC: ventral nerve cord. Scale bars: 500 μm. (H-J’, L-O) and 100 μm (P).

Altogether, these results indicate that grafts of the anterior and posterior planarian tips, corresponding to the intersections between the DVb and the midine, which contain *notum*- and *wnt1*-expressing cells, respectively, can induce the formation of a patterned head and tail, whereas grafts of the DVb or the midline alone cannot.

### Activation of Wnt/β-catenin signaling during regeneration promotes stem cell proliferation

Although the anterior and posterior tips can induce growth when grafted, our data and previous studies show that DVb grafts are also sufficient to promote growth (**Fig 1F**)^40,43^. Since the planarian tips also contain a portion of the DVb, *notum* and *wnt1* expression at the tips might solely pattern the forming outgrowth without affecting proliferation.

To test this, we analyzed whether these morphogens are required for neoblast proliferation during normal regeneration. Inhibition of *notum* and *wnt1* during regeneration can produce a shift in polarity^20,22^. However, under our RNAi conditions, the most frequent phenotypes were headless animals after *notum* inhibition and tailless animals after *wnt1* inhibition, which failed to regenerate the head or the tail, respectively, and displayed U-shaped VNCs at the tips (**S2A-F Fig**), as previously described^20,22^. We hypothesized that these phenotypes might reflect not only a failure to re-establish anterior and posterior identities, but also a reduced proliferative response of neoblasts. Our results show that inhibition of either *notum* or *wnt1* significantly decreased the number of phosphohistone-3^+^ (pH3^+^) cells throughout the whole-body at 6 hours of regeneration (hR), which corresponds to the early, body-wide peak of proliferation^46^ (**S2G Fig and Table S1**). When anterior and posterior wounds were analyzed separately at the same timepoint, *wnt1* inhibition reduced pH3^+^ cells at both wounds, whereas *notum* inhibition reduced it only at anterior wounds (**Fig 2A-D and Table S1**). The number of pH3^+^ cells were not affected in *wnt1* nor *notum* RNAi animals during the second wound-localized peak at 48hR^46^ neither in the whole-body data (**S2H Fig and Table S1**), nor locally in anterior or posterior wounds (**Fig 2B, 2D and Table S1**). These results indicate that *notum* and *wnt1* are required for the early and transient body-wide proliferative response of neoblasts during regeneration, with the effect of *notum* restricted to anterior wounds and that of *wnt1* extending to both anterior and posterior wounds.

**Figure 2.**
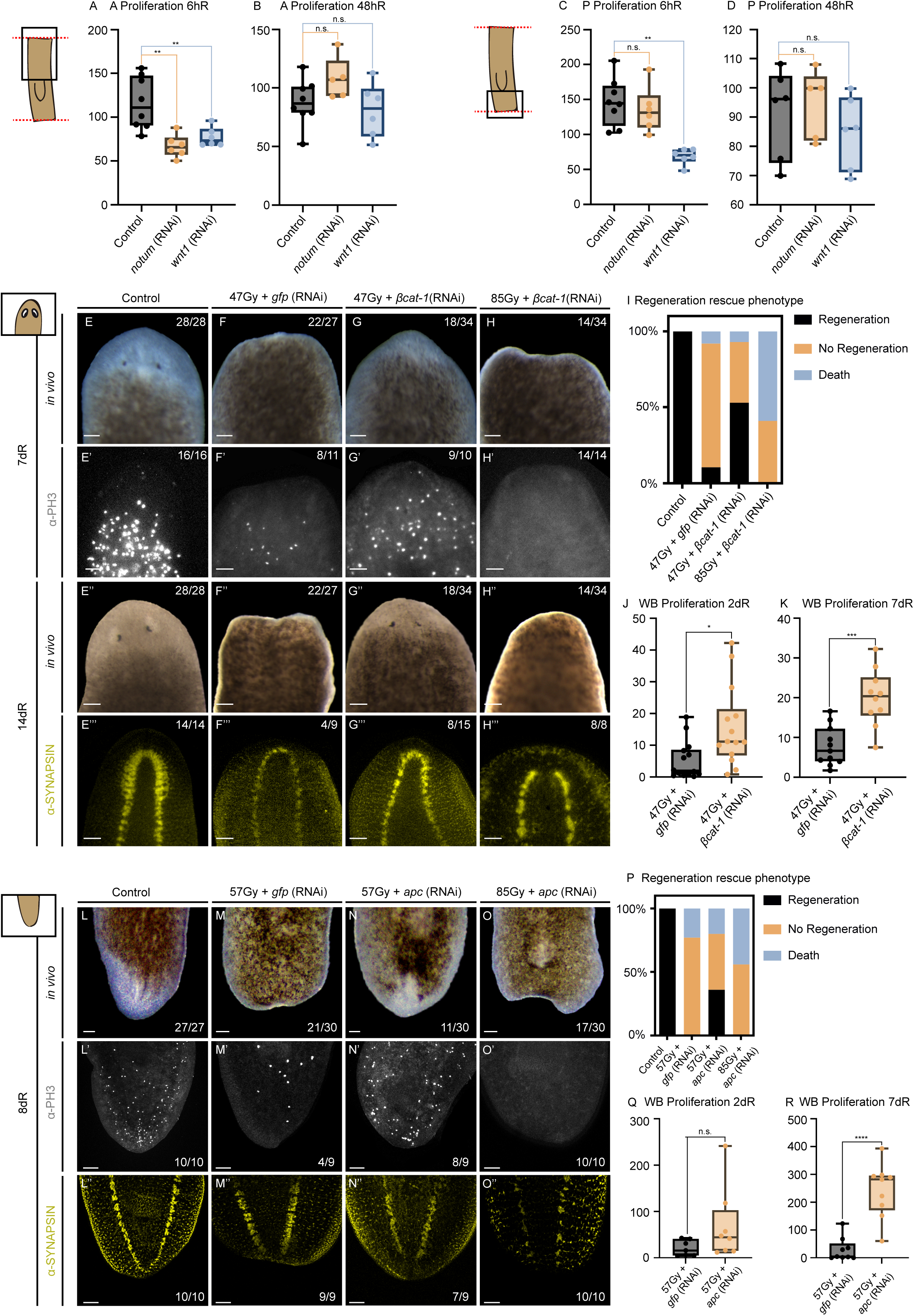
Activation of Wnt/β-catenin signaling during regeneration promotes stem cell proliferation. A-D. Quantification of neoblast proliferation (PH3^+^ cells/area) in the anterior (A-B) and posterior (C-D) regions of control, *notum* (RNAi) and *wnt1* (RNAi) planarians at 6hR (A, C) and 48hR (B, D). At 6hR, inhibition of both *notum* and *wnt1* reduces anterior proliferation (A), whereas only *wnt1* inhibition reduces posterior proliferation (C). Neither *notum* nor *wnt1* inhibition affects proliferation at 48hR (B, D). Statistical analysis: one-way ANOVA; **p < 0.01. **E-I.** Rescue of anterior regeneration in non-irradiated controls, sublethally irradiated planarians injected with *gfp* or *βcatenin-1* dsRNA, and lethally irradiated planarians injected with *βcatenin-1* dsRNA. **E-H.** *In vivo* phenotype at 7dR. **E’-H’.** α-PH3 immunostaining (grey) at 7dR, showing PH3^+^ cells in anterior blastemas of sublethally irradiated *βcatenin-1* (RNAi) planarians (G’) compared with *gfp* (RNAi) planarians (F’). **E’’-H’’.** *In vivo* phenotype at 14dR, showing the regeneration of a head with eyes in sublethally irradiated *βcatenin-1* (RNAi) planarians (G’’). **E’’’-H’’’.** α-synapsin immunostaining (yellow) at 14dR. Non-irradiated controls (E’’’) and sublethally irradiated *βcatenin-1* (RNAi) planarians (G’’’) regenerate the brain, sublethally irradiated *gfp* (RNAi) planarians show U-shaped VNCs (headless phenotype) (F’’’), and lethally irradiated *βcatenin-1* (RNAi) planarians fail to regenerate the brain (H’’’). **I.** Stacked bar plot showing the proportion of each phenotype in the conditions shown in E-H. **J-K.** Quantification of whole-body proliferation (PH3^+^ cells/area) in sublethally irradiated *gfp* (RNAi) and *βcatenin-1* (RNAi) planarians at 2dR (J) and 7dR (K), showing a significant increase after *βcatenin-1* inhibition at both time points. Statistical analysis: t-test; *p < 0.05 (J), ***p < 0.001 (K). **L-P.** Rescue of posterior regeneration in non-irradiated controls, sublethally irradiated planarians injected with *gfp* or *apc* dsRNA, and lethally irradiated planarians injected with *apc* dsRNA. **L-O.** *In vivo* phenotype at 8dR. **L’-O’.** α-PH3 immunostaining (grey) at 8dR, showing PH3^+^ cells in posterior blastemas of sublethally irradiated *apc* (RNAi) planarians (N’) compared with *gfp* (RNAi) planarians (M’). **L’’-O’’.** α-synapsin immunostaining (yellow) at 8dR. Non-irradiated controls (L’’) and sublethally irradiated *apc* (RNAi) planarians (N’’) regenerate the VNCs, whereas sublethally irradiated *gfp* (RNAi) (M’’) and lethally irradiated *apc* (RNAi) (O’’) planarians do not. **P.** Stacked bar plot showing the proportion of each phenotype in the conditions shown in L-O. **Q-R.** Quantification of whole-body proliferation (PH3^+^ cells/area) in sublethally irradiated *gfp* (RNAi) and *apc* (RNAi) planarians at 2dR (Q) and 8dR (R), showing an increase after *apc* inhibition that is significant only at 8dR. Statistical analysis: t-test; ****p < 0.0001 (R). The anterior and posterior regions correspond to the areas between the anterior end of the pharynx and the anterior-facing wound, and between the posterior end of the pharynx and the posterior-facing wound, respectively. In the rescue experiments, planarians were injected 3h before amputation and 3h, 6h and 24h after amputation (S2Q Fig). dR: days of regeneration. hR: hours of regeneration. VNC: ventral nerve cord. WB: whole body. Scale bars: 500 μm.

To further study how cWnt signaling regulates neoblast proliferation, we took advantage of the sensitivity of neoblasts to irradiation^25,47^. After establishing 85 Gy as the lethal dose in our planarian culture, we identified sublethal doses that impaired regeneration without causing death (**S2I-P Fig**). Our aim was to test whether downregulation or upregulation of the cWnt pathway in sublethally irradiated animals, through inhibition of *βcatenin-1* or *apc* (a component of the destruction complex^48^), respectively, could activate the proliferation of the surviving neoblasts and rescue regeneration. While lethally irradiated animals could not regenerate any structure (**S2K-K’’** and **S2N-N’’ Fig**), sublethally irradiated planarians displayed a phenotype reminiscent of *notum* and *wnt1* inhibition. Thus, 6/13 anterior and 5/12 posterior regenerating animals showed U-shaped anterior or posterior VNCs, corresponding to headless and tailless phenotypes (**S2J-J’, S2M-M’ and S2O-P Fig**), and lacked *notum* or *wnt1* expression at the anterior and posterior tips, respectively (**S2J’’ and S2M’’ Fig**). The dose producing the highest proportion of headless and tailless animals differed between anterior and posterior regeneration, being 47 Gy for anterior (**S2J-J’’ Fig**) and 57 Gy for posterior regeneration (**S2M-M’’ Fig**). Those doses were used for the rescue experiments. Irradiated animals were amputated, and *βcatenin-1* or *apc* was inhibited by dsRNA injections around the time of amputation and during the first 24h of regeneration, thus restricting the perturbation to the onset of the regenerative response (**Fig 2E-R and S2Q Fig**).

In the anterior-facing wounds rescue experiment, all non-irradiated control planarians regenerated (**Fig 2E-E’’’ and 2I**), while only 3/27 sublethally irradiated *gfp* (RNAi) animals did so (**Fig 2F-F’’’ and 2I**). Inhibition of *βcatenin-1* rescued anterior regeneration in 18/34 sublethally irradiated animals (**Fig 2G-G’’’ and 2I**), which regenerated a thinner, brain compared with non-irradiated controls (**Fig 2E’’’ and 2G’’’**). By contrast, *βcatenin-1* inhibition did not rescue regeneration in lethally irradiated animals (**Fig 2H-H’’’ and 2I**), indicating that the rescue depends on the neoblasts that remain after sublethal irradiation. Consistently, neoblast proliferation was significantly increased in sublethally irradiated *βcatenin-1* (RNAi) animals compared with sublethally irradiated *gfp* (RNAi) animals at 2 days of regeneration (dR), one day after the last dsRNA injection, and at 7dR, both in the whole body and in the anterior region (**Fig 2G’, 2J-K, S2R Fig and Table S1**).

For posterior regeneration, we inhibited *apc* in posterior-regenerating planarians irradiated at the sublethal and lethal doses (**Fig 2L-P**). None of the sublethally irradiated *gfp* (RNAi) animals regenerated the tail (**Fig 2M-M’’ and 2P**), probably because the sublethal dose used for posterior regeneration was higher than that used for anterior regeneration (57 Gy vs 47 Gy). In contrast, tail regeneration was rescued in 11/30 sublethally irradiated *apc* (RNAi) animals (**Fig 2N-N’’ and 2P**), and 7/9 of the regenerated tails showed VNCs resembling those of non-irradiated controls (**Fig 2L’’ and 2N’’**). None of the lethally irradiated *apc* (RNAi) animals regenerated (**Fig 2O-O’’ and 2P**), indicating that regeneration in sublethally irradiated animals relies on the remaining neoblasts. Neoblast proliferation in sublethally irradiated *apc* (RNAi) animals showed a non-significant increase at 2dR and was significantly increased at 8dR compared with sublethally irradiated *gfp* (RNAi) animals, both in the whole body and in the posterior region (**Fig 2N’, 2Q-R, S2S Fig, and Table S1**).

Interestingly, sublethally irradiated *βcatenin-1* (RNAi) animals also regenerated at posterior wounds, where 12/34 formed a head instead of a tail (**S2T-T’’’ Fig**). Similarly, sublethally irradiated *apc* (RNAi) animals regenerated at anterior wounds, giving rise to cycloptic animals in 5/30 cases (**S2U-U’’’ Fig**). These observations agree with the increase in proliferation observed in both conditions, which was not restricted to anterior or posterior wounds **(Fig 2J-K, 2Q-R, S2V-W Fig and Table S1**).

Finally, we explored whether cWnt signaling could influence the regulation of proliferation-related genes at the onset of regeneration. To this end, we analyzed the ATAC-seq and H3K27ac ChIP-seq datasets at 12hR previously generated by our group^49^. Most posterior-specific and anterior-specific enhancers close after *wnt1* and *notum* inhibition (75.5% and 87.7%, respectively; **S2X Fig**). An unbiased motif enrichment analysis of the CREs associated with these cWnt-dependent enhancers revealed, among others, binding motifs for transcription factors involved in cell proliferation. These included FOXM1 and MYB3R1 in posterior enhancers and NF-Y and AP-1 family members (Fra1, Fra2, JunB) in anterior enhancers (**S2Y-Z Fig and Table S2**). This suggests that cWnt-dependent chromatin remodeling at the wounds could contribute to the regulation of neoblast proliferation.

Altogether, these data indicate that cWnt signaling is required for the early proliferative response of neoblasts during regeneration and that either inhibition or activation of the pathway increases the proliferation of the remaining neoblasts in sublethally irradiated animals, rescuing regeneration of any wound.

### Wnt1 is required for the differentiation of ChAT neurons and parenchymal cells in posterior blastemas

The formation of a patterned nervous system in the outgrowths induced by tip grafts (**Fig 1J and 1N**) suggests that the signals present at the tips could influence not only neoblast proliferation but also cell differentiation. cWnt signaling is known to be required for brain patterning and for the differentiation of specific neuronal subtypes during anterior regeneration^50,51^. However, whether cWnt signaling is also required for tissue differentiation in posterior blastemas remains unknown (**Fig 3A**).

**Figure 3.**
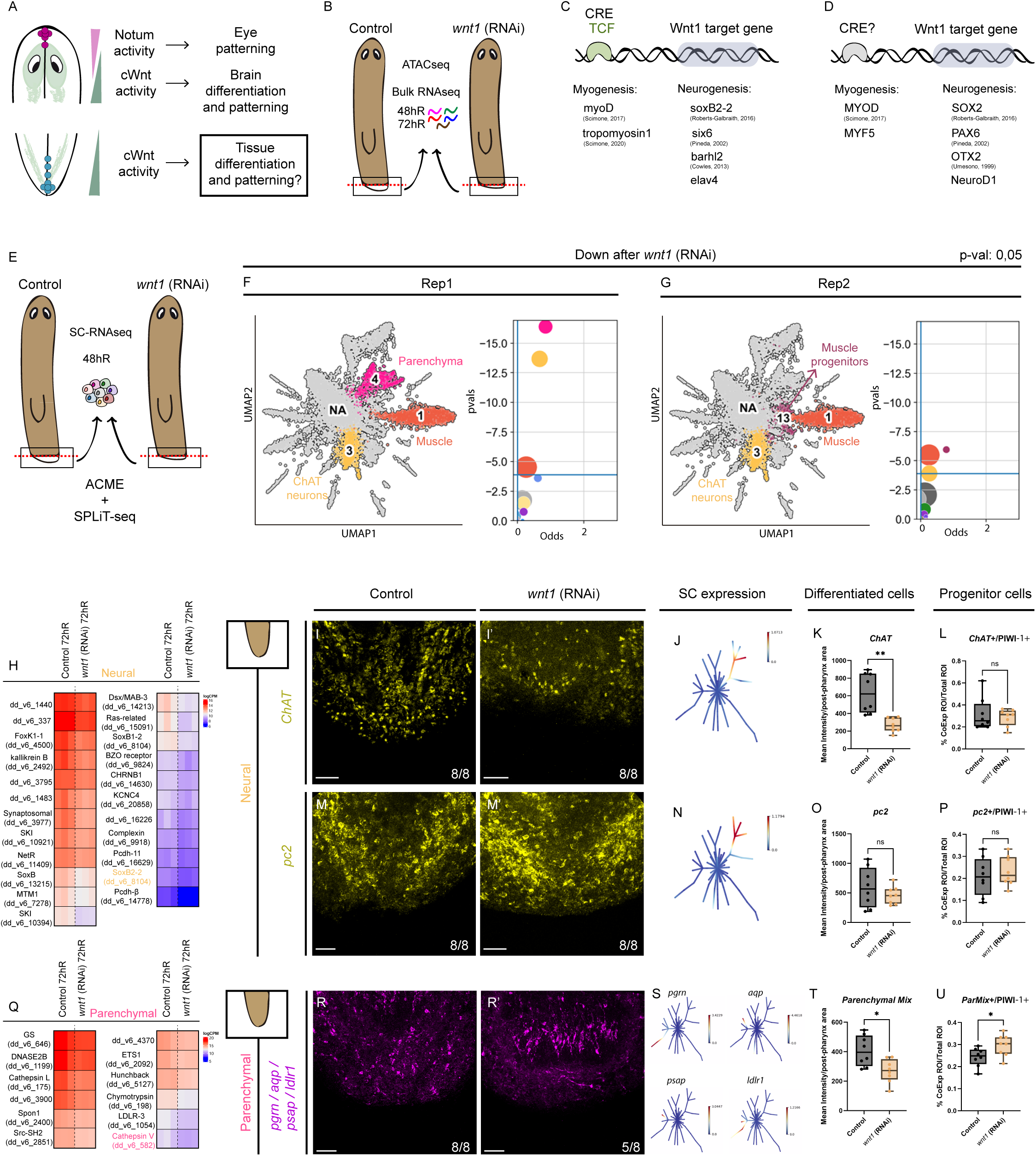
Wnt1 is required for the differentiation of ChAT neurons and parenchymal cells in posterior blastemas. **A.** Schematic summary of the known roles of Wnt/β-catenin signaling in tissue morphogenesis in planarians. **B.** Schematic representation of the collection of posterior blastemas of control and *wnt1* (RNAi) planarians for ATAC-seq and bulk RNA-seq^49^. **C.** Genes downregulated after *wnt1* inhibition that contain TCF binding motifs in their CREs, including genes involved in myogenesis (*myoD* and *tropomyosin-1*) and neurogenesis (*soxB2-2*, *six6*, *barhl2* and *elav4*). **D.** Unbiased motif enrichment analysis of the CREs of genes downregulated after *wnt1* inhibition, showing binding motifs of TFs involved in myogenesis (MyoD and Myf5) and neurogenesis (Sox2, Pax6, Otx2 and NeuroD1). **E.** Schematic representation of the collection of posterior blastemas of control and *wnt1* (RNAi) planarians at 48hR for scRNA-seq (ACME and SPLiT-seq). **F-G.** Comparison of cluster composition between control and *wnt1* (RNAi) samples in replicate 1 (F) and replicate 2 (G). UMAPs highlight the significantly reduced clusters, and plots show the odds ratio and p-value of each cluster (p < 0.05). The muscle body and ChAT neuron clusters are reduced in both replicates, whereas the parenchymal and muscle progenitor clusters are reduced in only one replicate. **H.** Heatmap of genes downregulated in *wnt1* (RNAi) posterior blastemas at 72hR (bulk RNA-seq) and expressed exclusively in neurons according to our scRNA-seq dataset. *soxB2-2*, whose inhibition produces indented tails (S3J-J’ Fig), is highlighted in yellow. **I-I’.** FISH showing *ChAT*^+^ cells (yellow) in posterior blastemas of control (I) and *wnt1* (RNAi) (I’) planarians at 48hR. **J.** *ChAT* expression in neural progenitors and differentiated ChAT neurons of intact planarians^53^. **K.** Quantification of *ChAT* signal intensity, showing a decrease after *wnt1* inhibition. **L.** Quantification of *ChAT*^+^/PIWI-1^+^ cells (ChAT progenitors), showing no significant differences after *wnt1* inhibition. **M-M’.** FISH showing *pc2*^+^ cells (yellow) in posterior blastemas of control (M) and *wnt1* (RNAi) (M’) planarians at 48hR. **N.** *pc2* expression in neural progenitors and differentiated neurons of intact planarians^53^. **O.** Quantification of *pc2* signal intensity, showing no significant differences after *wnt1* inhibition. **P.** Quantification of *pc2*^+^/PIWI-1^+^ cells (neural progenitors), showing no significant differences after *wnt1* inhibition. **Q.** Heatmap of genes downregulated in *wnt1* (RNAi) posterior blastemas at 72hR (bulk RNA-seq) and expressed exclusively in parenchymal cells according to our scRNA-seq dataset. *cathepsin V*, whose inhibition produces smaller tails (S3T-T’ Fig), is highlighted in pink. **R-R’.** FISH with a mix of probes against parenchymal markers (*pgrn*, *aqp*, *psap* and *ldlr1*; parenchymal mix, magenta) in posterior blastemas of control (R) and *wnt1* (RNAi) (R’) planarians at 48hR. **S.** Expression of *pgrn*, *aqp*, *psap* and *ldlr1* in parenchymal cells of intact planarians^53^. **T.** Quantification of the parenchymal mix signal intensity, showing a decrease after *wnt1* inhibition. **U.** Quantification of parenchymal mix^+^/PIWI-1^+^ cells (parenchymal progenitors), showing an increase after *wnt1* inhibition. Signal intensities (K, O, T) are normalized to the posterior region, between the posterior end of the pharynx and the posterior-facing wound, and co-expressing cells (L, P, U) to the total number of cells. Statistical analysis: t-test; **p < 0.01 (K), *p < 0.05 (T-U). All RNAi planarians were injected for two consecutive rounds, with head and tail amputation at the end of each round (Methods). CRE: *cis*-regulatory element. FISH: whole-mount fluorescent *in situ* hybridization. hR: hours of regeneration. TF: transcription factor. Scale bars: 500 μm.

As a first approach, we asked whether genes involved in tissue differentiation could be targets of Wnt1 in posterior blastemas. To this end, we integrated the ATAC-seq and bulk RNA-seq datasets of control and *wnt1* (RNAi) posterior blastemas previously generated by our group^49^ (**Fig 3B**). Among the genes downregulated after *wnt1* inhibition, those containing TCF binding motifs in their CREs included genes related to myogenesis, such as *myoD*^28^ and *tropomyosin-1*, and to neurogenesis, such as *soxB2-2*^36^, *six6*, *barhl2* and *elav4* (**Fig 3C and Table S2**). Accordingly, the GO terms associated with these genes included muscle organ development, regulation of muscle contraction, neurogenesis and synapse organization (**S3A Fig and Table S2**). In addition, an unbiased motif enrichment analysis of the CREs associated with genes downregulated after *wnt1* inhibition revealed binding motifs for transcription factors involved in myogenesis, such as MyoD and Myf5, and in neurogenesis, such as Sox2, Pax6, Otx2 and NeuroD1 (**Fig 3D and Table S2**). Together, these data suggest that Wnt1 could regulate the expression of genes involved in muscle and neural differentiation during posterior regeneration.

To determine which cell types are affected by *wnt1* inhibition, we generated a single-cell RNA-seq (scRNA-seq) dataset of posterior blastemas from control (*gfp* (RNAi)) and *wnt1* (RNAi) animals at 48hR, when the differentiation of new tissues is known to begin^41^, using ACME dissociation and SPLiT-seq^52^ (**Fig 3E, S4 Fig and Table S3**). The resulting UMAP recovered the major cell types present in planarian tails, whereas no clusters corresponding to eyes or pharynx were detected, consistent with the posterior origin of the samples (**S3B Fig**). We generated two replicates per condition and compared the proportion of cells assigned to each cluster between control and *wnt1* (RNAi) samples in each replicate. The muscle and ChAT neuron clusters were significantly reduced in *wnt1* (RNAi) animals in both replicates, while the parenchymal and muscle progenitor clusters were significantly reduced in only one replicate each (**p < 0.05; Fig 3F-G**). Moreover, the clusters showing a lower number of transcripts in *wnt1* (RNAi) samples relative to controls also included ChAT neurons, parenchyma, muscle body, muscle progenitors and secretory cells 1 and 4 (**S3C-H Fig**).

To validate these observations, we combined the scRNA-seq data with the bulk RNA-seq data of control and *wnt1* (RNAi) posterior blastemas at 72hR^49^ (**Fig 3B and 3E**). We selected the genes downregulated after *wnt1* inhibition whose expression was restricted to neural, parenchymal, muscle or secretory cells according to both our scRNA-seq dataset and the planarian cell type atlas^53^ (**Table S4**). The neural genes downregulated after *wnt1* inhibition included genes involved in synapse formation and *sox* genes, which are known to promote the differentiation of neural progenitors in planarians^54,55^ (**Fig 3H and Table S4**). Consistent with a role in posterior regeneration, inhibition of one of these genes, *soxB2-2*, produced indented tails (**S3I-K Fig**). Since our scRNA-seq data indicated that ChAT neurons were the only neural cluster reduced after *wnt1* inhibition, we next analyzed differentiated ChAT neurons (*chat*^+^ cells) and ChAT progenitors (*chat*^+^/PIWI-1^+^ cells) in *wnt1* (RNAi) blastemas (**Fig 3I-L**). The intensity of the *chat* signal was decreased in *wnt1* (RNAi) animals compared with controls (**Fig 3I-I’, 3K and Table S5**), while the proportion of *chat*^+^/PIWI-1^+^ cells was not affected (**Fig 3L, S3L-O’’’ Fig and Table S6**), suggesting that Wnt1 could be required for the terminal differentiation of ChAT neurons. In contrast, neither the signal intensity of the pan-neural marker *pc2* (**Fig 3M-O and Table S5**) nor the proportion of *pc2*^+^/PIWI-1^+^ cells (**Fig 3P, S3P-S’’’ Fig and Table S6**) was altered in *wnt1* (RNAi) animals. This suggests that Wnt1 is not generally required for neural differentiation but rather for the differentiation of specific neuronal subtypes, in agreement with the scRNA-seq data.

The parenchymal cluster was also reduced in one of the scRNA-seq replicates, and several parenchyma-specific genes, such as *ldlr-3*, *chymotrypsin* and *cathepsin V*, were downregulated in the bulk RNA-seq of *wnt1* (RNAi) blastemas (**Fig 3Q and Table S4**). Consistent with a role in posterior regeneration, inhibition of *cathepsin V* produced smaller tails (**S3T-U Fig**). To analyze the role of Wnt1 in parenchymal differentiation, we quantified differentiated parenchymal cells using a combination of probes against genes expressed in the four main parenchymal clusters (*pgrn, aqp, psap* and *ldlr1*)^53^ (**Fig 3R-S**), together with PIWI-1 immunostaining to identify parenchymal progenitors. The intensity of the *pgrn/aqp/psap/ldlr1* signal was reduced in *wnt1* (RNAi) animals (**Fig 3R-R’, 3T and Table S5**), while the proportion of cells co-expressing PIWI-1 was increased (**Fig 3U, S3V-Y’’’ Fig and Table S6**), suggesting that Wnt1 is required for the terminal differentiation of parenchymal cells.

Finally, although no secretory cluster was significantly reduced in our scRNA-seq dataset, the secretory 1 and 4 cluster showed the second strongest decrease in the number of expressed genes in *wnt1* (RNAi) samples (**S3H Fig**). In addition, 11 secretory-specific genes were downregulated in the bulk RNA-seq of *wnt1* (RNAi) blastemas (**Table S4**). These included hippocalcin and calmodulin, involved in calcium signaling, and *peptidase inhibitor 16*, whose inhibition produced duplicated tails (**S3Z-Z’’ Fig**).

Altogether, these results suggest that Wnt1 is required during posterior regeneration for the terminal differentiation of specific cell types, including ChAT neurons and parenchymal cells.

### Wnt/β-catenin signaling is necessary for muscle patterning and differentiation

The muscle cluster was reduced in *wnt1* (RNAi) samples in both scRNA-seq replicates (**Fig 3F-G**), and the muscle progenitor genes *myoD* and *nkx1-1*^28^ were downregulated after *wnt1* inhibition in the bulk RNA-seq (**Fig 4A and Table S4**). We therefore analyzed the differentiation and organization of muscle cells in *wnt1* (RNAi) animals.

**Figure 4.**
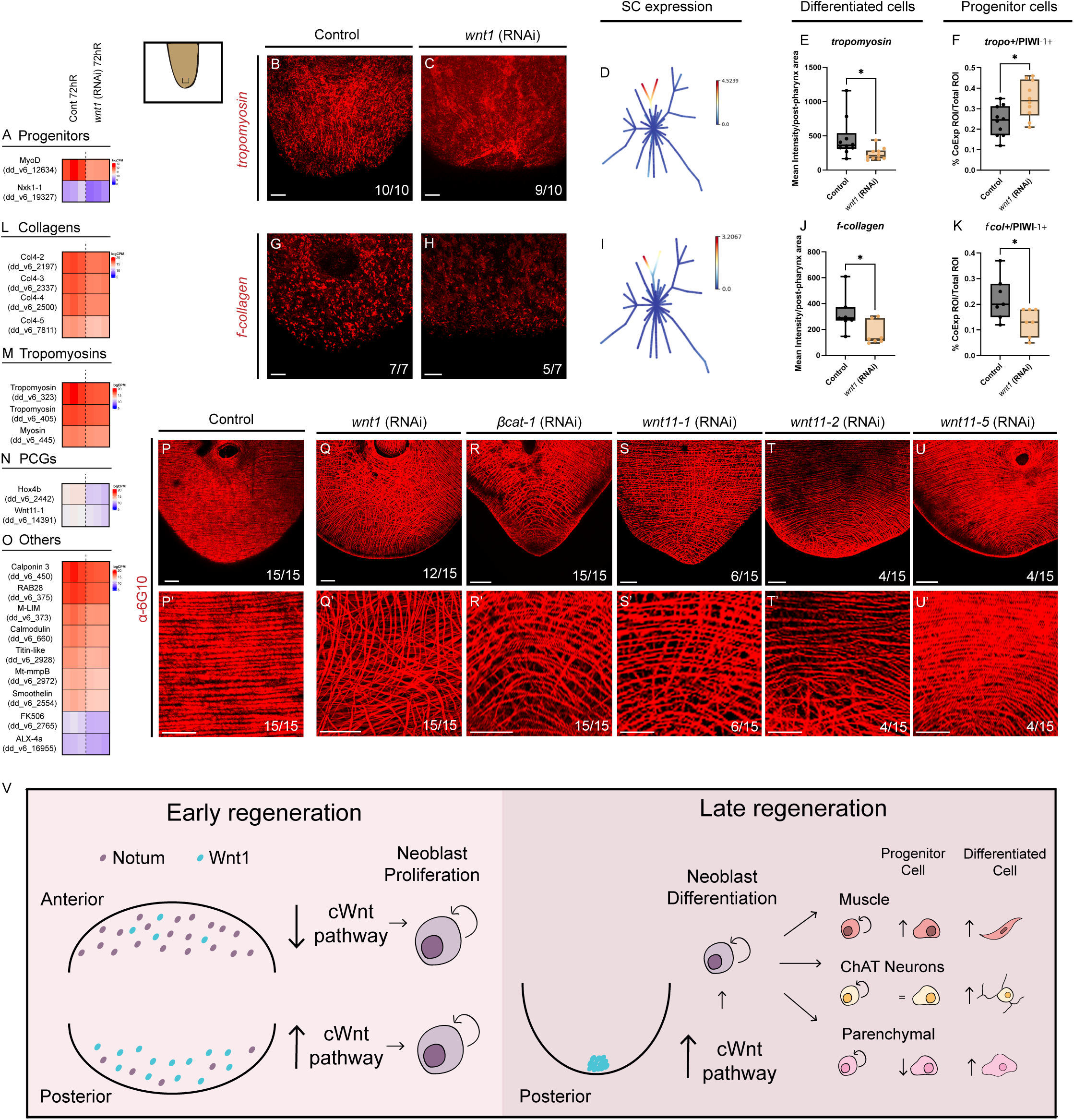
cWnt signaling regulates muscle differentiation and fiber organization during regeneration. **A.** Heatmap of genes downregulated in *wnt1* (RNAi) posterior blastemas at 72hR (bulk RNA-seq) and expressed in muscle progenitors according to our scRNA-seq dataset. **B-C.** FISH showing *tropomyosin* (*tropo*)^+^ cells (red) in posterior blastemas of control (B) and *wnt1* (RNAi) (C) planarians at 48hR. **D.** *tropo* expression in muscle progenitors and differentiated muscle of intact planarians^53^. **E.** Quantification of *tropo* signal intensity, showing a decrease after *wnt1* inhibition. **F.** Quantification of *tropo*^+^/PIWI-1^+^ cells (muscle progenitors), showing an increase after *wnt1* inhibition. **G-H.** FISH showing *f-collagen* (*fcol*)^+^ cells (red) in posterior blastemas of control (G) and *wnt1* (RNAi) (H) planarians at 48hR. **I.** *fcol* expression in committed muscle progenitors and differentiated muscle of intact planarians^53^. **J.** Quantification of *fcol* signal intensity, showing a decrease after *wnt1* inhibition. **K.** Quantification of *fcol*^+^/PIWI-1^+^ cells (committed muscle progenitors), showing a decrease after *wnt1* inhibition. Signal intensities (E, J) are normalized to the posterior region, between the posterior end of the pharynx and the posterior-facing wound, and co-expressing cells (F, K) to the total number of cells. Statistical analysis: t-test; *p < 0.05. **L-O.** Heatmaps of genes downregulated in *wnt1* (RNAi) posterior blastemas at 72hR (bulk RNA-seq) and expressed in muscle cells according to our scRNA-seq dataset, grouped into collagens (L), tropomyosins (M), PCGs (N) and others (O). **P-U’.** α-6G10 immunostaining (red) of posterior blastemas of control, *wnt1* (RNAi), *βcatenin-1* (RNAi), *wnt11-1* (RNAi), *wnt11-2* (RNAi) and *wnt11-5* (RNAi) planarians at 7dR. P’-U’ are magnifications of the midline region of the posterior tip. **V.** Graphical summary. During early regeneration, the inhibition of cWnt signaling by Notum at anterior wounds and its activation by Wnt1 at posterior wounds are required for neoblast proliferation. During late regeneration, Wnt1 at the posterior pole is required for the differentiation of committed muscle progenitors and muscle cells, ChAT neurons and parenchymal cells. All RNAi planarians were injected for two consecutive rounds, with head and tail amputation at the end of each round (Methods). dR: days of regeneration. FISH: whole-mount fluorescent *in situ* hybridization. hR: hours of regeneration. PCG: position control gene. Scale bars: 500 μm (B-C, G-H and P-U) and 50 μm (P’-U’).

To assess muscle differentiation in *wnt1* (RNAi) posterior blastemas, we quantified the signal intensity of the general muscle marker *tropomyosin-1* (*tropo*) (**Fig 4B-E**) and the proportion of *tropo*^+^ cells co-expressing PIWI-1. The *tropo* signal intensity was decreased in *wnt1* (RNAi) animals (**Fig 4B-C, 4E and Table S5**), while the proportion of *tropo*^+^/PIWI-1^+^ cells was increased (**Fig 4F, S5A-D’’’ Fig and Table S6**), suggesting that Wnt1 is involved in the final steps of muscle differentiation. To further test this hypothesis, we analyzed *f-collagen* (*fcol*), a marker of terminally differentiated muscle cells^56,57^ (**Fig 4G-I**). Both the *fcol* signal intensity and the proportion of *fcol*^+^/PIWI-1^+^ cells were decreased in *wnt1* (RNAi) animals (**Fig 4G-K, S5E-H’’’ Fig, Table S5 and Table S6**). Together with the scRNA-seq data, these results suggest that, after *wnt1* inhibition, muscle progenitors accumulate at an early stage (*tropo*^+^/PIWI-1^+^) and fail to progress towards later stages of differentiation, leading to a reduction of differentiated muscle. The decrease in *fcol*^+^/PIWI-1^+^ cells, which likely correspond to more advanced muscle progenitors^53,57^ (**Fig 4I**), could also explain the reduction of muscle progenitors observed in one of the scRNA-seq replicates (**Fig 3G**) and the downregulation of *myoD* and *nkx1-1* (**Fig 4A**). Consistent with a reduction of muscle cells, several muscle-related genes were downregulated in the bulk RNA-seq of *wnt1* (RNAi) blastemas, including different collagens and tropomyosins, calmodulin and titin-like (**Fig 4L-O and Table S4**). The muscle-expressed posterior PCGs *hox4b* and *wnt11-1* were also downregulated (**Fig 4N and Table S4**), although this may also reflect the failure to re-establish posterior identity after *wnt1* inhibition.

We next analyzed the organization of the muscle fibers after inhibition of Wnt pathway components required for posterior regeneration, namely *wnt1*, *βcatenin-1*, *wnt11-1*, *wnt11-2* and *wnt11-5*, using the muscle-specific antibodies 6G1058 and TMUS59. Inhibition of *wnt1* and *βcatenin-1* produced a general disorganization of the muscle fibers in posterior blastemas (12/15 and 15/15 animals, respectively; **Fig 4P-R’**), which in *wnt1* (RNAi) animals was also detected with the TMUS antibody (**S5I-J’ Fig**). A similar disorganization was observed in 6/15 *wnt11-1* (RNAi) animals (**Fig 4S-S’**), while inhibition of *wnt11-2* and *wnt11-5* produced a milder disorganization restricted to the tail tip in 4/15 animals each (**Fig 4T-U’**). In contrast, inhibition of the non-canonical Wnts wnt5 and wnt11-4 did not affect the organization of the muscle fibers in posterior blastemas (**S5K-L’ Fig**).

Since cWnt signaling is required for brain patterning and differentiation at anterior wounds^50,51^, we next asked whether it also regulates muscle differentiation in anterior blastemas. Inhibition of *wnt1* or *notum* altered neither the signal intensity of *tropo* and *fcol* nor the proportion of *tropo*^+^/PIWI-1^+^ and *fcol*^+^/PIWI-1^+^ cells in anterior blastemas (**S5M-AD Fig, Table S5 and Table S6**). This suggests that neither gene is required for muscle differentiation at anterior wounds, and that the requirement of *wnt1* for muscle differentiation is specific to posterior blastemas. However, *notum* inhibition disorganized the muscle fibers at the anterior tip (5/10 and 6/10 animals with the 6G10 and TMUS antibodies, respectively; **S5AE-AH’ Fig**), suggesting that *notum* contributes to the organization of the newly regenerated muscle. *βcatenin-1* inhibition also disorganized the muscle fibers at the anterior tip in 5/15 animals (**S5AI-AI’ Fig**). Since both the increase (*notum* RNAi) and the decrease (*βcatenin-1* RNAi) of cWnt signaling disrupted anterior fiber organization, these results suggest that a fine tune regulation of cWnt signaling levels is also required for the organization of the regenerating muscle at anterior wounds.

Together, these results suggest that cWnt signaling at posterior wounds is required for the terminal differentiation of muscle cells and for the organization of the regenerating muscle fibers, and that *notum* contributes to muscle fiber organization at anterior wounds.

## Discussion

In this work, we addressed two questions: whether the planarian tips act as organizers, and whether, as described for organizers in animals that regenerate specific structures, they regulate tissue growth and differentiation beyond axial patterning. Our results show that the planarian tips can trigger the formation of patterned heads and tails when grafted. They also show that the role of the cWnt signaling is not only defining the identity of these tips but also to regulate neoblast proliferation and tissue differentiation.

A defining property of organizers is their ability to induce the patterned growth of adjacent tissues^1–4,60^. Here, we show that, upon grafting, the anterior and posterior planarian tips can induce the formation of a patterned head and tail, respectively. Recently, grafting of the DVb combined with *βcatenin-1* inhibition was shown to produce head-like outgrowths with anterior identity^43^. This approach is comparable to our grafts of the planarian tips, which contain DVb signals together with regulators of cWnt signaling (Notum or Wnt1), suggesting that the combination of both types of signals, DVb and cWNT regulation, is required to produce bilateral outgrowths. Notably, the same principle seems to apply to both tips, since regions with opposite cWnt activity, low at the Notum-expressing anterior tip and high at the Wnt1-expressing posterior tip, can both produce bilateral structures, with anterior or posterior identity, respectively. Although we did not directly assess the origin of the cells forming our outgrowths, previous grafting studies that distinguished donor and host contributions, including grafts of the anterior tip, have shown that the outgrowths induced by planarian grafts are formed by host cells^40,44^. Together, these results support that the planarian tips have organizing capacity. However, since DVb grafts alone are sufficient to induce outgrowths (**Fig 1F**)^40,43^, Notum and Wnt1 at the tips could be only patterning the outgrowth without contributing to its growth.

In fact, inhibition of BMP/ADMP pathway components such as *bmp*, *admp* or *smad1* impairs regeneration^38,39,42^, suggesting that DVb signals could be responsible for stimulating the proliferation of host neoblasts. We therefore asked whether, in addition to BMP/ADMP signaling, the cWnt pathway also contributes to promote neoblast proliferation. Our results show that both *notum* and *wnt1* are required for the early, body-wide peak of neoblast proliferation at 6hR^46^. In agreement with this, Wnt1 was previously shown to signal through the Fzd1 and LRP5/6 receptors not only to specify posterior fate but also to trigger a proliferative response during posterior regeneration^61^. At this time point, *wnt1* is expressed equally in a scattered pattern at both anterior and posterior wounds, whereas *notum* is preferentially expressed at anterior wounds^20,22^. This could explain why *wnt1* inhibition reduces proliferation at both wounds, while *notum* inhibition only does so anteriorly. Moreover, neither gene affected the second, wound-localized peak of proliferation at 48hR, suggesting that, once their expression becomes restricted to the poles, *notum* and *wnt1* no longer regulate neoblast proliferation. Thus, besides patterning, *notum* and *wnt1* have a role in the early and transient activation of neoblast proliferation during regeneration.

Previous studies have shown that inhibition of *βcatenin-1* restores head regeneration in planarian species with limited regenerative abilities^62–65^, but they did not analyze whether neoblast proliferation increases when regeneration is rescued. To further study the role of cWnt signaling in neoblast proliferation, we therefore tested whether inhibition of *βcatenin-1* or *apc* could rescue regeneration in sublethally irradiated *S. mediterranea*. Both treatments partially rescued regeneration and increased the proliferation of the remaining neoblasts, showing that both downregulation and upregulation of cWnt signaling can enhance neoblast proliferation under these conditions. The absence of rescue in lethally irradiated animals, in which all neoblasts are depleted, further indicates that the rescue depends on the remaining neoblasts.

Interestingly, sublethally irradiated planarians displayed a phenotype similar to *notum* and *wnt1* (RNAi) animals, with U-shaped VNCs and headless or tailless phenotypes at anterior and posterior wounds, respectively. Some planarian species that cannot regenerate the head in nature, such as *Procotyla fluviatilis* and *Dendrocoelum lacteum*, also show a headless phenotype after head amputation, and their head regeneration can be restored by silencing *βcatenin-1*^62,64^. Together, these observations suggest that the modulation of cWnt signaling is required not only to specify head or tail identity but also to promote neoblast proliferation and trigger a growth response (**Fig 4V**). Notably, this ability does not depend on the type of wound. *βcatenin-1* inhibition in sublethally irradiated animals produced heads at both anterior and posterior wounds, and *apc* inhibition also rescued regeneration at anterior wounds. The mechanism by which cWnt signaling stimulates neoblast proliferation remains to be elucidated. A previous study reported that βcatenin-1 mediates head regrowth after DNA damage^66^, but its molecular interactions with neoblasts remain unexplored. In this regard, the binding motifs for proliferation-related transcription factors, such as FOXM1 and NF-Y, found in cWnt-dependent enhancers at 12hR (**S2Y-Z Fig**) point to candidate regulators that could link cWnt signaling to neoblast proliferation.

Regeneration requires not only the proliferation of stem cells but the coordination of cell proliferation, patterning and differentiation^67^. Having established the involvement of cWnt signaling in neoblast proliferation, we therefore focused on its role in cell differentiation. cWnt signaling had been previously implicated in the differentiation of specific neuronal subtypes in the regenerating brain^50,51^, but its role in the differentiation of posterior tissues had been less studied and remained unknown. Our transcriptomic and histological data suggest that *wnt1* is required for the terminal differentiation of ChAT neurons, muscle cells and parenchymal cells (**Fig 4V**). Planarian ChAT neurons are a widely distributed neuronal subpopulation present in the central nervous system (brain and VNCs) and the eyes, and they have been implicated in the control of muscle function, since their inhibition produces motor defects^68,69^. Thus, the decrease in ChAT neurons could be explained by a direct role of Wnt1 in the differentiation of ChAT neurons or it could be related to the decrease in muscle cells observed in *wnt1* (RNAi) animals. Wnt1 is also required for the organization of the regenerating muscle fibers in posterior blastemas. Similar defects were observed after inhibition of other Wnt pathway components required for posterior regeneration, such as *βcatenin-1*, *wnt11-1*, *wnt11-2* and *wnt11-5*, as well as after inhibition of *notum* and *βcatenin-1* at anterior wounds. These organization defects could be secondary to an altered innervation by ChAT neurons or reflect a direct effect on muscle, since several Wnts are expressed in planarian muscle cells^21,28^ and cWnt signaling regulates muscle differentiation as shown in this study and in several embryonic and regenerative contexts^70^. Parenchymal cells are a relatively unexplored planarian cell population that includes phagocytic cells and glia^35,36,71^. Among their few known functions are the support and maintenance of neural populations and the promotion of neural regeneration^36^. It is therefore tempting to speculate that the differentiation of ChAT neurons and parenchymal cells could share cWnt-dependent regulatory elements. Finally, the defects observed in the differentiation of muscular and neural cells are unlikely to result from a reduced proliferative response of neoblasts. Neoblast proliferation at posterior wounds was not affected by *wnt1* inhibition at 48hR, and the progenitors of these cell types were not reduced. Nevertheless, we cannot exclude that the failure to re-establish posterior identity after *wnt1* inhibition also contributes to the defects seen in cell differentiation.

We also found that several genes expressed in secretory cells, particularly in the secretory 4 population^53^, are downregulated after *wnt1* inhibition. However, no secretory cluster was significantly reduced in our scRNA-seq data. This could be due to the small size of these clusters, or to the fact that secretory 4 cells are grouped with secretory 1 cells in our dataset, which could mask a specific reduction. Alternatively, *wnt1* inhibition might affect the expression of specific genes in secretory 4 cells rather than their number. The lack of well-defined markers for planarian secretory cells limited further analysis in our study. A study on the logic of planarian neoblast differentiation suggests that neural, secretory and muscle cells share a common differentiation path^72^. Since these three cell types are affected after *wnt1* inhibition, cWnt signaling could act at the beginning of this path. However, several observations argue against an exclusive role at its onset. ChAT progenitors were not reduced and early muscle progenitors accumulated, the reduction of neurons was restricted to ChAT neurons, and the evidence for a decrease in secretory cells is weak. Together, these data suggest that cWnt signaling acts at later steps of this path, possibly with distinct roles in each lineage.

The control of multiple processes through signaling pathways such as the cWnt pathway is a feature of organizing regions extensively documented in embryonic and regenerative model systems^73–78^. Our data provides experimental evidence that, in planarians, cWnt signaling regulates proliferation and differentiation in addition to its well-established role in axial patterning. Further studies will be required to understand how cWnt signaling controls the entry of neoblasts into mitosis and orchestrates their differentiation to regenerate the missing structures in coordination with pre-existing tissues.

Finally, returning to our initial questions, our results support that the planarian tips have organizing capacity, as they can trigger the formation of patterned heads and tails when grafted. Moreover, the cWnt signaling that define these tips is not restricted to axial patterning. It also contributes to the early proliferative response of neoblasts and to the terminal differentiation of specific cell types, functions classically attributed to the organizers of animals that regenerate specific structures. Our results together with the pre-exiting data suggest that the early wound-induced cWnt signaling that precedes pole formation could be responsible to trigger the proliferative response, since when modulating the Wnt signal we only observe an effect in the early proliferative peak, while the late cWnt signal restricted to the poles could be the one responsible of cell-identity specification and differentiation. Favoring this hypothesis, inhibition of pole-related genes as *islet-1*, impairs specifically the late *wnt1* signal and produces tailless/bi-headed animals^79,80^.

Altogether, our findings suggest that coupling growth and differentiation to patterning may be a common feature of regenerative organizers, regardless of whether regeneration involves a single structure or the whole body.

## Methods

### Planarian maintenance

The planarian species used in this study was *Schmidtea mediterranea*, clonal strain BCN-10. Animals were maintained in glass containers and transferred to Petri dishes for experimental procedures. Both during maintenance and experiments, planarians were kept at 20 °C in planarian artificial medium (PAM) containing 0.016 mM NaCl (Merck, 7647145), 0.01 mM MgSO4·7H2O (Merck, 105886), 0.012 mM NaHCO3 (Merck, 106329), 0.001 mM KCl (Merck, 104936), 0.001 mM MgCl2·6H2O (Merck, 105833) and 0.01 mM CaCl2·2H2O (Merck, 102382) in Milli-Q water. Animals were fed organic beef liver twice per week and were starved for 1 week after any experimental procedure. Animals of approximately 7 mm in length were used for most experiments, and animals of approximately 15 mm for grafting experiments.

### Grafting

Grafts were performed as previously described^45^. Two 15-mm planarians (host and donor) were placed on a wet square of cigarette paper on an ice-chilled glass slide. The graft insertion site was punched out from the dorsal surface of the immobilized host by pressing a square capillary tube (0.6 mm exterior diameter) vertically, and the tissue plug was removed with a micro spatula and a clean brush. The graft was then resected from the donor animal using a square capillary (0.8 mm interior diameter), collected with a micro spatula, placed on top of the insertion site and gently pushed down with a small, clean brush until fully inserted. Host animals were then decapitated. To improve graft retention, hosts were tightly wrapped in cigarette paper, transferred to a black Petri dish (to prevent photosensitive responses) containing 10 ml of chilled planarian water (4 °C), and kept at 4 °C for 4 h. Finally, the cigarette paper was removed, and the planarians were allowed to recover at 20 °C. Anterior tips were grafted into the post-pharyngeal region and posterior tips into the pre-pharyngeal region of the host. DVb tissue from the lateral part of the head and midline tissue from the pre-pharyngeal region were transplanted into the post-pharyngeal region.

### RNAi

Double-stranded RNA (dsRNA) for *wnt1*, *notum*, *βcatenin-1*, *apc*, *wnt11-1*, *wnt11-2*, *wnt11-4*, *wnt11-5* and *wnt5* was synthesized as previously described^81^. dsRNA (1 μg/μl) was administered with a Nanoject II (Drummond Scientific, Broomall, PA, United States) as three injections of 32 nl per day. Unless otherwise indicated, the injection protocol consisted of two rounds of injections on 3 consecutive days, and 1 day after the last injection of each round, planarians were amputated pre- and post-pharyngeally to induce anterior and posterior regeneration. Control animals were injected with dsRNA against *gfp*, a gene absent in planarians.

### Irradiation and regeneration rescue experiments

Planarians were irradiated with a RX Smart de Yxlon Int at 47 Gy or 57 Gy, which were established as sublethal doses for anterior and posterior regeneration, respectively, or at 85 Gy, which was established as the lethal dose. Three days after irradiation, animals were amputated to induce anterior or posterior regeneration and injected with *βcatenin-1*, *apc* or *gfp* dsRNA 3 h before amputation and 3 h, 6 h and 24 h after amputation. Animals were analyzed *in vivo* and fixed at different time points after the amputation.

### Immunostaining

Whole-mount immunostaining was performed as previously described^82^. Animals were killed with cold 2% HCl and fixed with 4% PFA at room temperature. After 4 h in blocking solution (1% BSA in PBS with 0.3% Triton X-100, PBSTx), animals were incubated overnight at 4 °C with primary antibodies, washed extensively with PBSTx, blocked for 2 h and incubated overnight at 4 °C with secondary antibodies. The following primary antibodies were used: mouse anti-synapsin (anti-SYNORF1, 3C11; 1:50; Developmental Studies Hybridoma Bank), anti-PH3 (1:1000; Developmental Studies Hybridoma Bank), anti-PIWI-1 (1:2000), anti-TMUS59 (1:1000) and anti-6G1058 (1:1000; Developmental Studies Hybridoma Bank). The secondary antibodies were Alexa 488-conjugated goat anti-mouse (1:400; Molecular Probes, A28175) and Alexa 568-conjugated goat anti-rabbit (1:1000; Molecular Probes, A-11011). Nuclei were stained with DAPI (1:5000), and samples were mounted in 70% glycerol in PBS.

### Fluorescent in situ hybridization (FISH)

FISH was performed as previously described^83,84^. DIG-labeled RNA probes were synthesized in vitro using T7 or SP6 RNA polymerases (Roche), purified by precipitation with ethanol and 7.5 M ammonium acetate, resuspended in 20 µl of ddH2O and adjusted to a final concentration of 200 ng/µl. Animals were killed in 7% N-acetyl-L-cysteine (NAC), fixed in 4% formaldehyde and stored in methanol at −20 °C. After overnight hybridization, samples were washed twice with 4× SSC containing Triton X-100 (SSCTx), three times with 0.2× SSCTx and with MABTween. Samples were then blocked in MABTween containing 5% horse serum and 0.5% Western blocking reagent (Roche) and incubated with anti-DIG-POD (1:2000; Roche, 11093274910). After washing the antibody for 2 h, the signal was developed with fluorescein-tyramide (1:500), 4-iodophenylboronic acid (4-IPBA; 1:1000) and H_2_O_2_ (1:15000). Samples were mounted in 70% glycerol in PBS.

### FISH combined with immunostaining

For the co-detection of tissue-specific transcripts and PIWI-1, animals were killed and fixed as described for FISH, and the FISH protocol was followed by the immunostaining protocol.

### ACME dissociation

Posterior blastemas from *gfp* (RNAi) and *wnt1* (RNAi) animals at 48hR were collected in two biological replicates per condition. Tissue dissociation and fixation were performed using ACME^52^ with the following modifications. Each sample, containing 30 posterior blastemas, was incubated for 5 min in 2 mL tubes and pipetted as in the original protocol to promote dissociation. Samples were filtered through 40 µm strainers (CellTrics) and centrifuged at 1000 × g for 5 min at 4 °C, and the pellet was resuspended in the remaining volume after discarding part of the supernatant. Samples were then filtered into 2 mL tubes using 1 mL filter tips (Flowmi). To remove the ACME solution, cells were washed with 1 mL of buffer (1× phosphate-buffered saline (PBS) with 1% bovine serum albumin (BSA)) and centrifuged at 1000 × g for 5 min at 4 °C. The supernatant was discarded, and cells were resuspended in 900 µL of buffer in 1.5 mL tubes, supplemented with 100 µL of DMSO and stored at −80 °C.

### SPLiT-seq

All samples were processed using the SPLiT-seq protocol as previously described^52^. Frozen dissociated cells (unsorted) were thawed, centrifuged twice at 1000 × g for 5 min at 4 °C, resuspended in ∼450 µL of buffer (1× PBS with 1% BSA) and filtered through a 50 µm strainer (CellTrics). For each sample, a 1:3 aliquot (50 µL of sample and 100 µL of buffer) was stained for 20 min at room temperature in the dark with 1.5 µL of DRAQ5 (0.5 mM stock) and 0.6 µL of Concanavalin-A conjugated with AlexaFluor 488 (1 mg/mL stock, Invitrogen), while the remaining cells were kept at 4 °C. To estimate cell concentration (singlets/µL), three measurements of 10 µL were taken for each aliquot by flow cytometry. Singlet events were counted using an FSC-H vs FSC-A gate to select singlets, a Concanavalin-A-positive gate to select cells with cytoplasm and a DRAQ5-positive gate to distinguish cells from cellular debris. The remaining unstained cells were diluted according to this singlet count in 0.5× PBS to a final working concentration of ∼400,000 cells/mL (3,000 singlets per well).

### Single-cell transcriptomic analysis

Single-cell RNA-seq data were analyzed using Python-based workflows. Cells were clustered using the Leiden algorithm^85^ at resolution 1 and visualized by UMAP, and clusters were annotated based on their marker genes and the planarian cell type atlas^53^. Differences in cluster composition between control and *wnt1* (RNAi) samples were assessed for each replicate considering p < 0.05 as significant. To compare transcriptional profiles between control and *wnt1* (RNAi) conditions at 48hR, we performed a pseudobulk analysis. Raw gene expression counts were aggregated within each cluster for each sample to generate pseudobulk expression matrices, an approach that minimizes single-cell technical noise while retaining cluster-specific biological variation. Differential gene expression analysis was then performed between control and *wnt1* (RNAi) pseudobulk samples within each cluster.

### Bulk RNA-seq analysis

Bulk RNA-seq data of control and *wnt1* (RNAi) posterior blastemas were obtained and analyzed as previously described^49^ (**Fig 3B**). Downregulated genes were manually filtered to retain those whose expression was restricted to neural, parenchymal, muscle or secretory cells according to our scRNA-seq dataset and the planarian cell type atlas^53^.

### ATAC-seq, ChIP-seq and motif analyses

ATAC-seq and H3K27ac ChIP-seq data from anterior and posterior blastemas at 12hR were obtained and analyzed as previously described^49^.

### Image acquisition and quantification

*In vivo* images were acquired with a Scmex 3.0 camera (DC.3000s, Visual Inspection Technology) mounted on a Zeiss Stemi SV 6 stereomicroscope. Brightfield colorimetric ISH and FISH images were obtained with a ProgRes C3 camera (Jenoptik, Jena, Germany). Confocal images of whole-mount immunostainings and FISH were acquired with a Zeiss LSM 880 confocal microscope (Zeiss, Oberkochen, Germany). Fiji/ImageJ^86^ was used to generate representative confocal projections for each experimental condition. FISH signal intensity was quantified in Fiji by measuring the mean signal intensity within the blastema and normalizing it to the blastema area (**Table S5**). PH3^+^ cells were counted and curated manually and normalized to the area of the analyzed region (whole body, anterior or posterior region).

### Co-expression quantification

Co-expression was quantified using a custom macro implemented in Fiji/ImageJ86. Multichannel images were first split into individual channels corresponding to DAPI, the tissue-specific marker (GenA) and PIWI-1 (GenB). The nuclear channel was processed by applying a Gaussian blur (σ = 2), followed by automatic thresholding using the Otsu method to generate a binary mask, and a watershed algorithm was applied to separate adjacent nuclei. To approximate a perinuclear region, the nuclear mask was expanded by iterative morphological dilation (three iterations, approximately one pixel per dilation). Particles were then detected from the dilated mask using size and circularity constraints, and the resulting regions of interest (ROIs) were added to the ROI Manager. These ROIs were used to measure the mean fluorescence intensity and integrated density of the GenA and GenB channels independently. Measurements from both channels were combined into a single dataset, with one row per ROI, and exported as a comma-separated values (CSV) file for downstream analysis. Mean fluorescence intensity values for each ROI were normalized per sample and per gene within an intensity range of 1,000-20,000 units, as values outside this range were not observed in single cells across the analyzed images. A ROI was classified as positive for a given channel when its intensity exceeded the mean intensity of that channel, and as co-expressing when it was positive for both GenA and GenB. The proportion of co-expressing ROIs was calculated by normalizing the number of double-positive ROIs to the total number of ROIs identified per sample.

### Statistical analysis

Statistical analyses were performed using GraphPad. Data normality was assessed with the Shapiro-Wilk test. Comparisons between two groups were performed using unpaired Student’s t-test, and comparisons among three groups (control, *notum* (RNAi) and *wnt1* (RNAi)) using one-way ANOVA. Significance levels are indicated in the figures as follows: ns, not significant; *p < 0.05; **p < 0.01; ***p < 0.001; ****p < 0.0001.

## Acknowledgments

We acknowledge all the members of the Adell-González lab, Cebrià lab and Solana Lab for comments and discussion.

