## Supplementary Figures 1_5 and legends for "cWnt signaling regulates the early proliferative response and the differentiation of the neural, muscle and parenchymal tissues during planarian whole-body regeneration"

**Supplementary Figures legends**

**S1 Fig. Grafting phenotypes. A.** Schematic representation of the grafting protocol (see Methods). **B.** Schematic representation of the anterior tip graft into the post-pharyngeal region of the host. **C.** *In vivo* "Polarized" anterior outgrowth 2 weeks after grafting. The red arrow marks an eye. **D.** FISH showing *pc2* (yellow) in a "Polarized" anterior outgrowth 2 weeks after grafting, showing an anteriorized localization of $pc2^{+}$ cells. Nuclei are labeled with DAPI (blue). **E.** Schematic representation of the posterior tip graft into the pre-pharyngeal region of the host. **F.** *In vivo* "Polarized-like" posterior outgrowth 2 weeks after grafting. **G.** FISH showing *pc2* (yellow) in a "Polarized-like" posterior outgrowth 2 weeks after grafting, showing scattered $pc2^{+}$ cells. FISH: whole-mount fluorescent *in situ* hybridization. VNC: ventral nerve cord. Scale bars: 500 μm (C-D, F-G).

**S2 Fig. Regeneration rescue phenotypes. A-F.** Control, *notum* (RNAi) and *wnt1* (RNAi) planarians at 10dR, *in vivo* (A-C) and after α-synapsin immunostaining (yellow) (D-F). *notum* and *wnt1* inhibition produce headless (B, E) and tailless (C, F) planarians, respectively, with VNCs fused in a U-shape (red arrows). **G-H.** Quantification of whole-body proliferation ($PH3^{+}$ cells/area) in control, *notum* (RNAi) and *wnt1* (RNAi) planarians at 6hR (G) and 48hR (H), showing a decrease exclusively at 6hR after *notum* and *wnt1* inhibition. Statistical analysis: one-way ANOVA; $^{**}p < 0.01$. **I-K''.** Non-irradiated, sublethally irradiated (47 Gy) and lethally irradiated (85 Gy) planarians at 8dR after head amputation, *in vivo* (I-K), after α-synapsin immunostaining (yellow) (I'-K') and after FISH for *notum* (magenta) (I''-K''). Irradiated planarians fail to regenerate anterior structures. Sublethally irradiated planarians show VNCs fused in a U-shape (J', red arrow), whereas lethally irradiated planarians fail to regenerate the VNCs (K'), and neither reestablishes *notum* expression. **L-N''.** Non-irradiated, sublethally irradiated (57 Gy) and lethally irradiated (85 Gy) planarians at 8dR after tail amputation, *in vivo* (L-N), after α-synapsin immunostaining (yellow) (L'-N') and after FISH for *wnt1* (cyan) (L''-N''). Irradiated planarians fail to regenerate the tail. Sublethally irradiated planarians show VNCs fused in a U-shape (M', red arrow), whereas lethally irradiated planarians fail to regenerate the VNCs (N'), and neither reestablishes *wnt1* expression. **O-P.** Stacked bar plots summarizing the nervous system phenotypes after head (O) and tail (P) amputation: normal brain or VNCs, smaller brain or VNCs, VNCs fused in a U-shape and cut VNCs. **Q.** Schematic representation of the regeneration rescue protocol. Planarians were irradiated 3 days before amputation and injected 3h before and 3h, 6h and 24h after amputation with 1 nl of dsRNA (2 μg/μl) per injection. **R-S.** Quantification of anterior proliferation in sublethally irradiated *gfp* (RNAi) and *βcatenin-1* (RNAi) planarians (R) and of posterior proliferation in sublethally irradiated *gfp* (RNAi) and *apc* (RNAi) planarians (S) ($PH3^{+}$ cells/area), showing an increase at 2dR and 7dR after *βcatenin-1* inhibition (R) and at 8dR after *apc* inhibition (S). **T-T'''.** *In vivo* posterior regeneration at 14dR of non-irradiated controls (T) and of sublethally irradiated planarians injected with *gfp* dsRNA, which fail to regenerate a complete tail (T'), sublethally irradiated planarians injected with *βcatenin-1* dsRNA, which regenerate a head instead of a tail (T''), and lethally irradiated planarians injected with *βcatenin-1* dsRNA, which fail to regenerate (T'''). **U-U'''.** *In vivo* anterior regeneration at 8dR of non-irradiated controls (U) and of sublethally irradiated planarians injected with *gfp* dsRNA, which fail to regenerate (U'), sublethally irradiated planarians injected with *apc* dsRNA, which regenerate cyclopic heads (U''), and lethally irradiated planarians injected with *apc* dsRNA, which fail to regenerate (U'''). **V-W.** Quantification of posterior proliferation in sublethally irradiated *gfp* (RNAi) and *βcatenin-1* (RNAi) planarians (V) and of anterior proliferation in sublethally irradiated *gfp* (RNAi) and *apc* (RNAi) planarians (W) ($PH3^{+}$ cells/area), showing an increase at 7dR after *βcatenin-1* inhibition (V) and at 8dR after *apc* inhibition (W). R-S and V-W. Statistical analysis: t-test; $^{*}p < 0.05$ (R, 2dR), $^{****}p < 0.0001$ (R, 7dR), $^{***}p < 0.001$ (S) and $^{**}p < 0.01$ (V-W). The anterior and posterior regions correspond to the areas between the anterior end of the pharynx and the anterior-facing wound, and between the posterior end of the pharynx and the posterior-facing wound, respectively. **X.** Schematic representation of the ATAC-seq and H3K27ac ChIP-seq data of anterior and posterior blastemas at 12hR[49], showing the percentage of posterior- and anterior-specific enhancers that close after *wnt1* and *notum* inhibition, respectively. **Y-Z.** Lollipop plots showing the TF binding motifs enriched in the CREs of posterior (Y) and anterior (Z) enhancers, including motifs of TFs related to cell proliferation. CRE: *cis*-regulatory element. dR: days of regeneration. FISH: whole-mount fluorescent *in situ* hybridization. hR: hours of regeneration. TF: transcription factor. VNC: ventral nerve cord. WB: whole body. Scale bars: 1 mm (A-F) and 500 µm (I-N'' and T-U''').

**S3 Fig. Tissue differentiation defects after *wnt1* inhibition. A.** Biological process GO terms of the genes downregulated after *wnt1* inhibition (bulk RNA-seq of posterior blastemas at 72hR) that contain TCF binding motifs in their CREs. GO terms related to myogenesis and neurogenesis are highlighted in red and yellow, respectively. **B.** UMAP of control and *wnt1* (RNAi) posterior blastemas at 48hR, showing the 27 clusters obtained with the Leiden algorithm at resolution 1. **C-G.** UMAPs of the clusters affected after *wnt1* inhibition, namely ChAT neurons (C, yellow), parenchyma (D, pink), muscle progenitors (E, red), muscle body (F, orange) and secretory 1, 4 (G, purple), each with a dot plot of its top 10 marker genes. Orange highlights two collagens also downregulated in the bulk RNA-seq of *wnt1* (RNAi) posterior blastemas at 72hR. **H.** Heatmap comparing the levels of gene expression per cluster between control (gfp_rep1 and gfp_rep2) and *wnt1* (RNAi) (wnt1_rep1 and wnt1_rep2) replicates, without statistical testing. **I-J'.** Posterior blastemas of control (I-I') and *soxB2-2* (RNAi) (J-J') planarians at 7dR, *in vivo* (I, J) and after α-synapsin immunostaining (yellow) (I', J'). *soxB2-2* inhibition produces indented tails with unfused VNCs. **K.** UMAP showing the enriched expression of *soxB2-2* in ChAT neurons. **L-S'''.** FISH combined with immunostaining showing *ChAT*$^+$ (L-O''') or *pc2*$^+$ (P-S''') cells (yellow) and PIWI-1$^+$ cells (green) in posterior blastemas of control (L-M''' and P-Q''') and *wnt1* (RNAi) (N-O''' and R-S''') planarians at 48hR. M-M''', O-O''', Q-Q''' and S-S''' are magnifications of L, N, P and R, respectively. Red arrows mark co-expressing cells. Nuclei are labeled with DAPI (blue). **T-T'.** Posterior blastemas of *cathepsin V* (*catV*) (RNAi) planarians at 7dR, *in vivo* (T) and after α-synapsin immunostaining (yellow) (T'), showing smaller blastemas and VNCs. **U.** UMAP showing the enriched expression of *cathepsin V* in parenchymal cells. **V-Y'''.** FISH combined with immunostaining showing parenchymal mix$^+$ (*psap*, *pgrn*, *aqp* and *ldlr1*) cells (magenta) and PIWI-1$^+$ cells (green) in posterior blastemas of control (V-W''') and *wnt1* (RNAi) (X-Y''') planarians at 48hR. W-W''' and Y-Y''' are magnifications of V and X, respectively. Red arrows mark co-expressing cells. Nuclei are labeled with DAPI (blue). **Z-Z'.** Posterior blastemas of *peptidase inhibitor 16* (*pep.inh.16*) (RNAi) planarians at 7dR, *in vivo* (Z) and after α-synapsin immunostaining (yellow) (Z'), showing double tails with duplicated VNCs. **Z''.** Heatmap of genes downregulated in *wnt1* (RNAi) posterior blastemas at 72hR (bulk RNA-seq) and expressed in secretory 1 and 4 cells according to our scRNA-seq dataset. *pep.inh.16* is highlighted in purple. All RNAi planarians were injected for two consecutive rounds, with head and tail amputation at the end of each round (Methods). CRE: *cis*-regulatory element. dR: days of regeneration. FISH: whole-mount fluorescent *in situ* hybridization. GO: Gene Ontology. hR: hours of regeneration. VNC: ventral nerve cord. Scale bars: 500 µm (I-J', T-T' and Z-Z') and 50 µm (L-S''' and V-Y''').

**S4 Fig. Quality control and clustering of the single-cell RNA-seq dataset of control and *wnt1* (RNAi) posterior blastemas at 48hR.** All panels correspond to the combined dataset of control (*gfp* (RNAi)) and *wnt1* (RNAi) samples (two biological replicates per condition), analyzed with Scanpy. **A-B.** Violin plots showing the number of genes detected per cell (n_genes_by_counts; A) and the total number of counts per cell (total_counts; B), on a logarithmic scale. **C-D.** Scatter plots of the number of genes detected versus the total number of counts per cell before (C) and after (D) quality-control filtering. **E-F.** Selection of highly variable genes, showing the normalized (E) and non-normalized (F) dispersion of each gene as a function of its mean expression, with highly variable genes in black and the remaining genes in grey. **G.** Variance ratio explained by each of the first 150 principal components, ranked in decreasing order and shown on a logarithmic scale, used to select the number of components for downstream analyses. **H-J.** Violin plots showing the number of genes (n_genes; H), the number of counts (n_counts; I) and the total counts (total_counts; J) per cell in each of the 27 clusters (0-26) obtained with the Leiden algorithm at resolution 1 (Leiden 1). **K-M.** UMAPs in which each cell is colored according to its number of genes (n_genes; K), number of counts (n_counts; L) and number of genes by counts (n_genes_by_counts; M). **N.** Dot plot showing the expression of the two top-ranked marker genes of each Leiden 1 cluster. Dot size represents the percentage of cells in each cluster expressing the gene, and color intensity represents its mean expression in that cluster. The dendrogram shows the hierarchical clustering of Leiden 1 clusters according to their expression profiles, which determines the order in which clusters are displayed.

**S5 Fig. cWnt signaling regulates muscle differentiation in posterior blastemas and muscle organization in anterior and posterior blastemas. A-H'''.** FISH combined with immunostaining showing *tropomyosin* (*tropo*)$^+$ (A-D''') or *f-collagen* (*fcol*)$^+$ (E-H''') cells (red) and PIWI-1$^+$ cells (green) in posterior blastemas of control (A-B''' and E-F''') and *wnt1* (RNAi) (C-D''' and G-H''') planarians at 48hR. B-B''', D-D''', F-F''' and H-H''' are magnifications of A, C, E and G, respectively. Yellow arrows mark co-expressing cells. Nuclei are labeled with DAPI (blue). **I-J'.** α-TMUS immunostaining (red) of posterior blastemas of control (I-I''') and *wnt1* (RNAi) (J-J') planarians at 7dR, showing the circular, diagonal and longitudinal muscle fibers of controls (I'-I''') and the disorganization of the regenerated muscle after *wnt1* inhibition (J-J'). **K-L'.** α-6G10 immunostaining (red) of posterior blastemas of *wnt11-4* (RNAi) (K-K') and *wnt5* (RNAi) (L-L') planarians at 7dR, showing well-organized regenerated muscle. The control is shown in Fig 4P-P'. I'-I''', J', K' and L' are magnifications of the midline region of the posterior tip. **M-X'''.** FISH combined with immunostaining showing *tropo*$^+$ (M-R''') or *fcol*$^+$ (S-X''') cells (red) and PIWI-1$^+$ cells (green) in anterior blastemas of control (M-N''' and S-T'''), *wnt1* (RNAi) (O-P''' and U-V''') and *notum* (RNAi) (Q-R''' and W-X''') planarians at 48hR. N-N''', P-P''', R-R''', T-T''', V-V''' and X-X''' are magnifications of M, O, Q, S, U and W, respectively. Yellow arrows mark co-expressing cells. Nuclei are labeled with DAPI (blue). **Y-Z''.** FISH showing *tropo*$^+$ (Y-Y'') and *fcol*$^+$ (Z-Z'') cells (red) in anterior blastemas of control (Y, Z), *wnt1* (RNAi) (Y', Z') and *notum* (RNAi) (Y'', Z'') planarians at 48hR. **AA-AD.** Quantification of *tropo* (AA) and *fcol* (AC) signal intensity and of *tropo*$^+$/PIWI-1$^+$ (AB) and *fcol*$^+$/PIWI-1$^+$ (AD) cells in anterior blastemas of control, *wnt1* (RNAi) and *notum* (RNAi) planarians, showing no significant differences. Signal intensities are normalized to the anterior region, between the anterior end of the pharynx and the anterior-facing wound, and co-expressing cells to the total number of cells. Statistical analysis: one-way ANOVA. **AE-AI'.** α-6G10 (AE, AG and AI) and α-TMUS (AF and AH) immunostaining (red) of anterior blastemas of control (AE-AF'), *notum* (RNAi) (AG-AH') and *βcatenin-1* (RNAi) (AI-AI') planarians at 7dR. Controls show well-organized regenerated muscle, whereas *notum* and *βcatenin-1* inhibition disorganizes the muscle at the anterior tip. AE', AF', AG', AH' and AI' are magnifications of the midline region of the anterior tip. dR: days of regeneration. FISH: whole-mount fluorescent *in situ* hybridization. hR: hours of regeneration. Scale bars: 500 μm (A, C, E, G, I, J, K, L, M, O, Q, S, U, W, Y-Z'', AE, AF, AG, AH and AI) and 50 μm (B-B''', D-D''', F-F''', H-H''', I'-I''', J', K', L', N-N''', P-P''', R-R''', T-T''', V-V''', X-X''', AE', AF', AG', AH' and AI').

### Supplementary figure 1

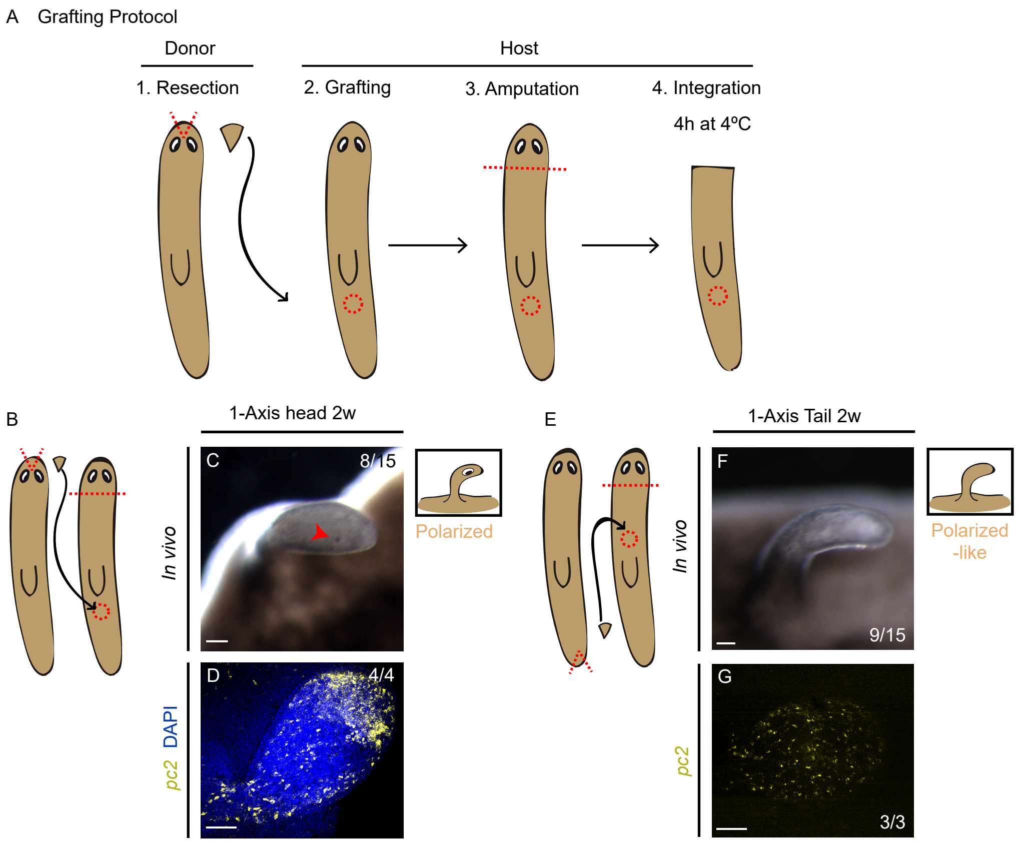

### Supplementary figure 2

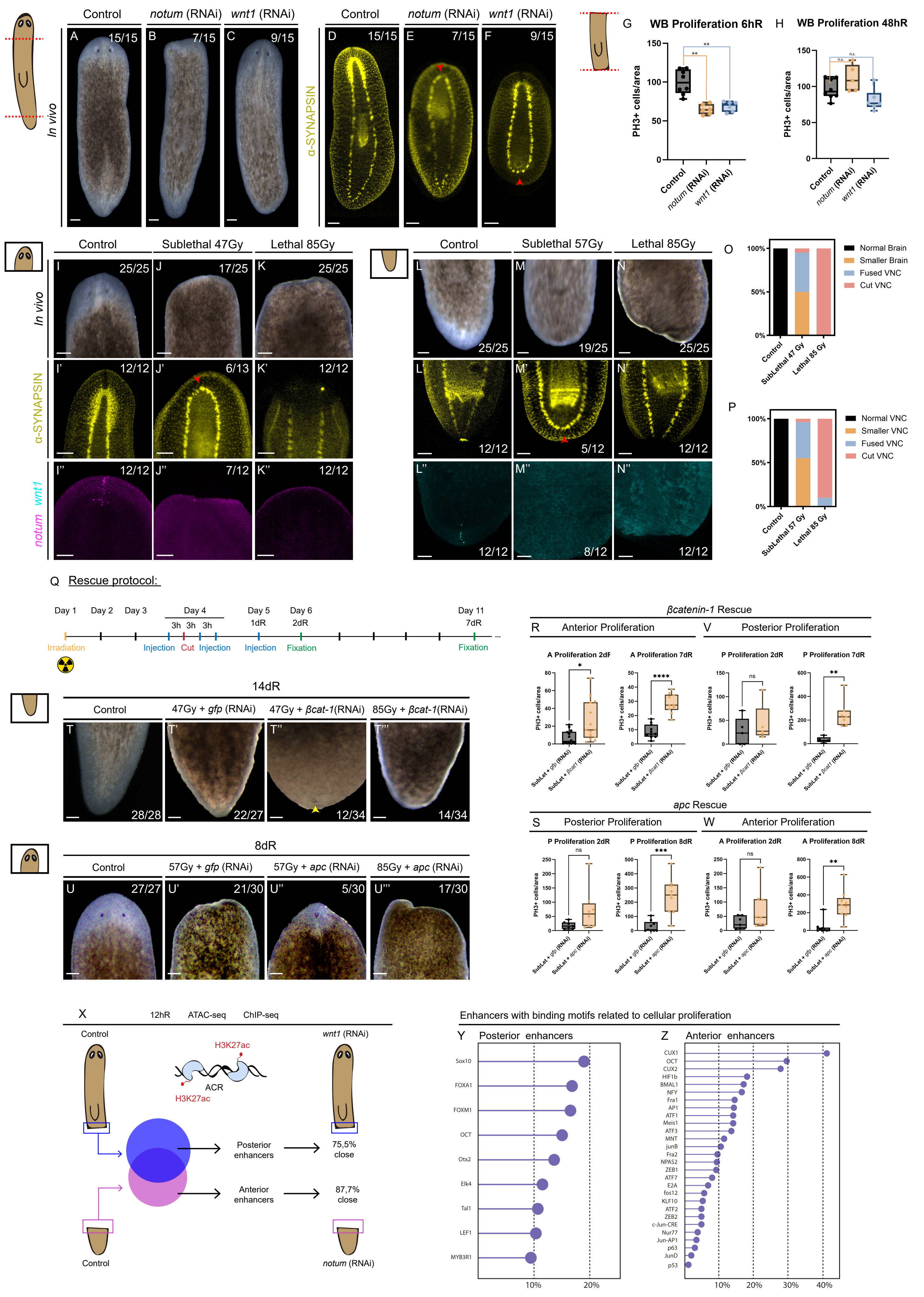

### Supplementary figure 3

A GOs of the down-regulated genes after wnt1 inhibition presenting a TCF binding motif in their CREs

| BIOLOGICAL PROCESS | | | | | |
|---|---|---|---|---|---|
| GO.ID | Term | Annotated | Significant | Expected | weightFisher |
| GO:0006355 | regulation of transcription, DNA-template | 573 | 18 | 05.04 | 6,00E-06 |
| GO:0038032 | termination of G protein-coupled receptor | 33 | 3 | 0.29 | 30 |
| GO:0016055 | Wnt signaling pathway | 36 | 3 | 0.32 | 38 |
| GO:0048749 | compound eye development | 1 | 1 | 0.01 | 88 |
| GO:0048730 | epidermis morphogenesis | 1 | 1 | 0.01 | 88 |
| GO:0071929 | alpha-tubulin acetylation | 1 | 1 | 0.01 | 88 |
| GO:0018160 | peptidyl-pyrromethane cofactor linkage | 2 | 1 | 0.02 | 175 |
| GO:0007517 | muscle organ development | 2 | 1 | 0.02 | 175 |
| GO:0006937 | regulation of muscle contraction | 2 | 1 | 0.02 | 175 |
| GO:0040034 | regulation of development, heterochronic | 2 | 1 | 0.02 | 175 |
| GO:0022008 | neurogenesis | 3 | 1 | 0.03 | 261 |
| GO:0007099 | centriole replication | 4 | 1 | 0.04 | 347 |
| GO:0000122 | negative regulation of transcription | 4 | 1 | 0.04 | 347 |
| GO:0048013 | ephrin receptor signaling pathway | 5 | 1 | 0.04 | 432 |
| GO:0050808 | synapse organization | 5 | 1 | 0.04 | 432 |
| GO:0009058 | biosynthetic process | 1800 | 26 | 15.82 | 439 |

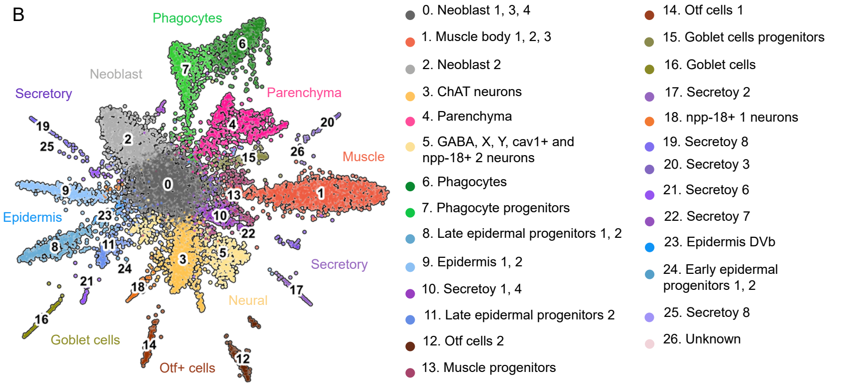


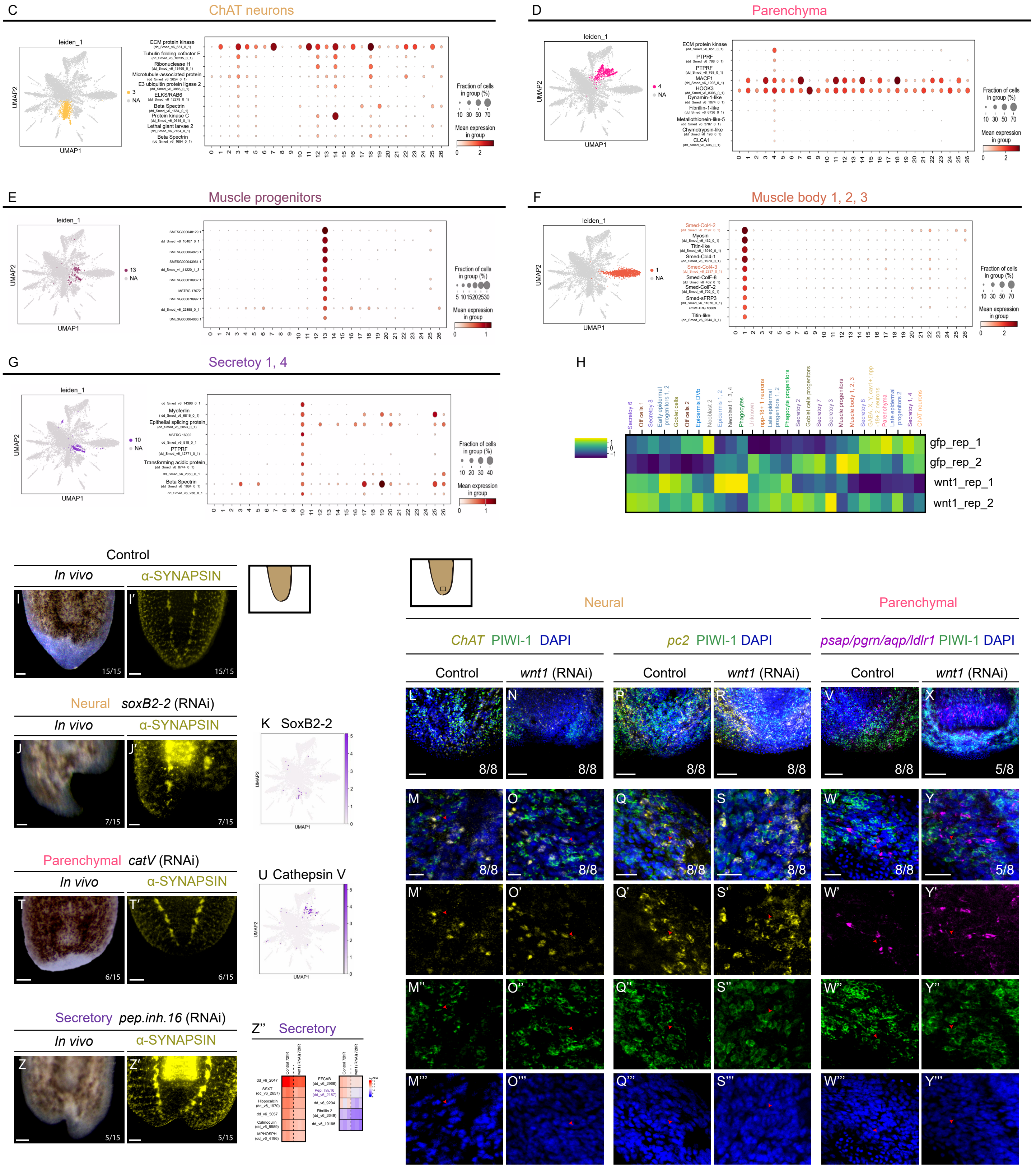

### Supplementary Figure 4

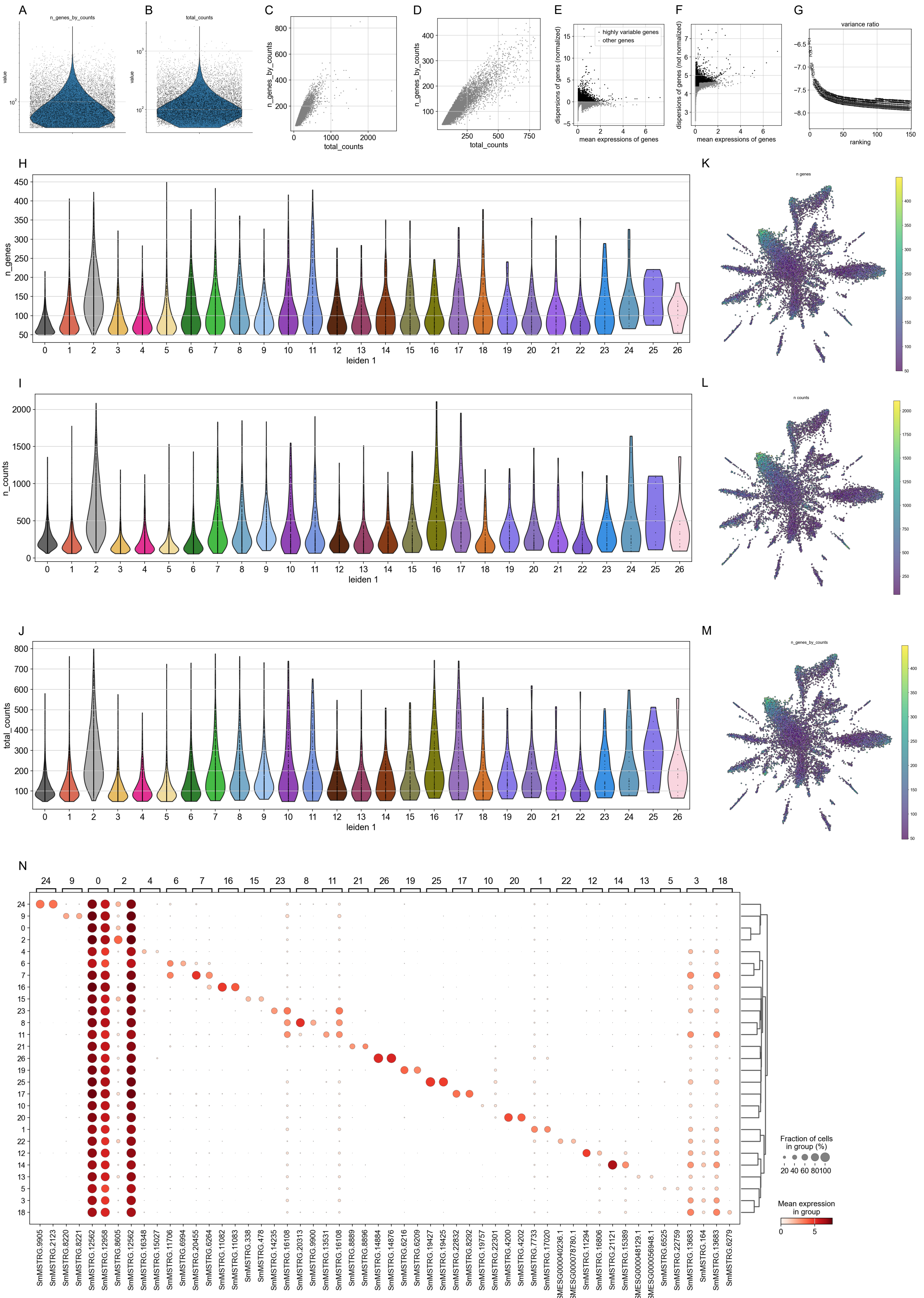

Supplementary figure 5

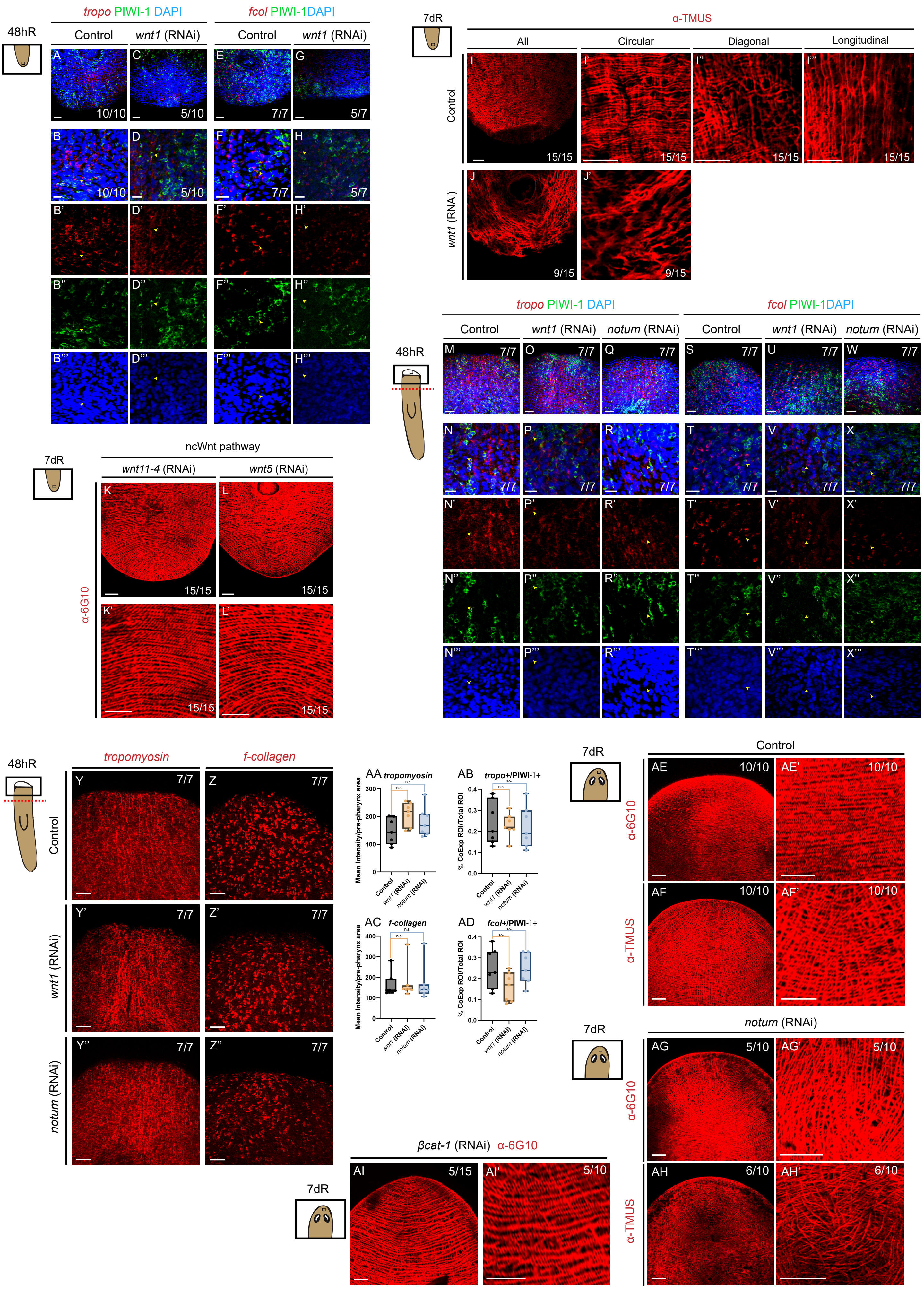

**Supplementary Tables legends**

**Table S1. Raw quantification of mitotic activity (PH3+ cells) underlying the proliferation analyses shown in Figure 2 and Figure S2**. Mitotic activity was quantified by counting all PH3+ cells in each planarian. That number was then normalized to the area of the corresponding region (anterior, posterior or whole body), measured on the same images, and values are expressed as PH3+ cells/mm². Data is shown for each individual planarian (Plan1, Plan2, etc.). Sheet S1 contains data for control, *notum* (RNAi) and *wnt1* (RNAi) animals. Sheet S2 contains the data for the rescue experiment with *β-catenin-1* (RNAi) performed after sublethal irradiation (47 Gy) . Sheet S3 contains the data for the rescue experiment with *apc* (RNAi) performed after sublethal irradiation (57 Gy). Details of the immunostaining, irradiation and quantification procedures are provided in Materials and Methods.

**Table S2. TCF binding motifs and transcription factor motif enrichment in cis-regulatory elements during early regeneration**. All analyses are based on the ATAC-seq and ChIP-seq datasets previously generated by our group (Pascual-Carreras et al., 2023). Sheets S1 and S2 list the genes containing TCF binding motifs in their cis-regulatory elements (CREs) in wild-type planarians at 12 hours of regeneration. For each gene, the genome identifier (Genome ID), the automatic annotation (Annotation), the TCF binding site motif identified (TCF binding site motif) and the gene name (Name) are shown. Genes with a described role in neurogenesis are highlighted in yellow and those with a described role in myogenesis are highlighted in red. Sheet S3 shows the Gene Ontology (GO) enrichment analysis of the genes listed in sheet S2 for the Biological Process and Molecular Function categories. For each GO term, the identifier (GO.ID), the term description (Term), the number of genes annotated with the term in the reference gene set (Annotated), the number of those genes present in the analysed gene list (Significant), the number expected by chance (Expected) and the p-value obtained with the weighted Fisher's exact test (weightFisher) are indicated. Sheets S4 and S5 show the unbiased motif enrichment analysis of all CREs associated with genes downregulated in wnt1(RNAi) anterior and posterior blastemas at 12 hours of regeneration. For each motif, the motif name (Motif Name), a short label (Label), the transcription factor family (Class) and the consensus sequence (Consensus) are given, together with the enrichment p-value (P-value), its logarithm (Log P-value) and the Benjamini–Hochberg-corrected q-value (q-value (Benjamini)), as well as the number and percentage of target and background sequences containing the motif. Motifs recognized by transcription factors with a described role in neurogenesis are highlighted in yellow and those with a described role in myogenesis are highlighted in red.

**Table S3. Single-cell RNA-seq analysis of control vs wnt1(RNAi) posterior blastemas at 48 hours of regeneration**. Sheet S1 lists the cell barcodes, the unique nucleotide sequences that tag all transcripts captured from an individual cell and thus identify each cell in the dataset, for the cells analysed from control (gfp (RNAi)) and wnt1(RNAi) posterior blastemas at 48 hours of regeneration, each with two biological replicates (replicates 1 and 2). Sheet S2 shows the number of cells assigned to each cluster obtained with the Leiden algorithm (Traag et al., 2019) at resolution 1 (Leiden 1) in each sample. Sheet S3 lists the marker genes identified for each Leiden 1 cluster. Sheet S4 contains the annotation coordinates used for read mapping and quantification, in which each transcript is placed at its genomic coordinates in the dd_Smes_g4 genome assembly (Grohme et al., 2018), as available in PlanMine (Brandl et al., 2016; Rozanski et al., 2019).

**Table S4. Differentially expressed genes in posterior blastemas of *wnt1* (RNAi) animals at 72 hours of regeneration**. Bulk RNA-seq comparison of posterior blastemas from control (RNAi) and wnt1(RNAi) planarians at 72 hours post-amputation. Genes were assigned to separate sheets after manual filtering of those whose expression in single-cell transcriptomic data was exclusive to neural (sheet S1), parenchymal (sheet S2), muscle (sheet S3) or secretory (sheet S4) cell populations. For each gene, the log2 fold change between wnt1(RNAi) and control blastemas (logFC) and the adjusted p-value (adj.P.Val) are shown, with positive and negative logFC values indicating up- and downregulation in wnt1(RNAi) animals, respectively. Each entry also includes the gene identifier in the genome annotation (Genome ID), the corresponding transcript in the dd_Smed_v6 transcriptome (dd_Smed_v6), the identifier used in the single-cell dataset (SC ID), the automatic annotation (Gene name) and the manually curated gene name (curated gene name). The columns "TCF presence in CREs (promoters)" and "TCF presence in CREs (enhancers)" indicate whether TCF binding motifs are present in the cis-regulatory elements associated with each gene, at promoters and enhancers respectively (Pascual-Carreras, 2023). The column "Down in βcat1 (RNAi) RNA-seq Tewari et al. (2019)" indicates whether the gene was downregulated upon β-catenin-1 RNAi in the posterior RNA-seq dataset reported by Tewari et al. (2019). Finally, "Intacts (planmine)" indicates the cell type in which each gene is expressed in the single-cell atlas of intact planarians (Plass et al., 2018) available through PlanMine (Rozanski et al., 2019), and "regeneration UMAP" indicates its expression in the single-cell dataset of regenerating planarians generated in this study.

**Table S5. Quantification of FISH signal intensity for neural, parenchymal and muscle markers in blastemas at 48 hours of regeneration**. Signal intensity of fluorescent *in situ* hybridization (FISH) staining was quantified in individual planarians at 48 hours of regeneration by measuring the area of the blastema and the signal intensity within that area using Fiji/ImageJ (Schindelin et al., 2012). Sheet S1 shows the quantification of the neural markers ChAT and pc2 in posterior blastemas of control and wnt1(RNAi) animals. Sheet S2 shows the same quantification for parenchymal cells, detected with a combination of probes against *psap*, *ldlr*, *aqp* and *pgrn*, in posterior blastemas of control and *wnt1* (RNAi) animals. Sheet S3 shows the quantification of the muscle markers *f-collagen* and *tropomyosin* in posterior blastemas of control and *wnt1* (RNAi) animals, as well as in anterior blastemas of control, *wnt1* (RNAi) and *notum* (RNAi) animals. Each sheet contains a separate table for each marker or probe combination, which includes a summary of the signal intensity normalized to blastema area for each condition, followed by the individual measurements of blastema area and signal intensity for each planarian.

**Table S6. Quantification of co-expression of tissue-specific markers with PIWI in posterior and anterior blastemas at 48 hours of regeneration**. Co-expression was quantified from FISH + immunostainings of planarians at 48 hours of regeneration using a custom Fiji/ImageJ macro (Schindelin et al., 2012), as described in Materials and Methods. Briefly, nuclei were segmented from the DAPI channel and expanded to include the perinuclear region, and each resulting region of interest (ROI) was considered an individual cell. A ROI was classified as positive for a given gene when its normalized mean fluorescence intensity exceeded the mean intensity of that channel in the sample, and as co-expressing when it was positive for both genes. Sheet S1 provides a summary of all the data. Sheets S2 to S8 show the quantification for each probe combination, namely ChAT and PIWI (sheet S2), pc2 and PIWI (sheet S3), the parenchymal probe mix (*psap, ldlr, aqp and pgrn*) and PIWI (sheet S4), *tropomyosin* and PIWI in posterior blastemas (sheet S5), *f-collagen* and PIWI in posterior blastemas (sheet S6), *tropomyosin* and PIWI in anterior blastemas (sheet S7) and *f-collagen* and PIWI in anterior blastemas (sheet S8). Sheets S2 to S6 include posterior blastemas of control (C1, C2, C3...) and wnt1(RNAi) (W1, W2, W3...) planarians, whereas sheets S7 and S8 include anterior blastemas of control, *wnt1* (RNAi) and *notum* (RNAi) (N1, N2, N3...) planarians. For each planarian, the table lists every ROI (ROI) with its normalized mean fluorescence intensity for the tissue-specific marker (Mean_GenA) and for *piwi* (Mean_GenB), whether it was classified as positive for each gene (PosA and PosB) and whether it showed co-expression (Coexpresion). An adjacent summary table indicates, for each planarian, the mean intensity of each channel used as the positivity threshold (MeanA and MeanB), the number of co-expressing ROIs (Total_CoExp), the total number of ROIs analysed (Total_Cells) and the proportion of co-expressing cells (CoExp/TotalCells).
